# Integrated analysis reveals neuro-immune pathway in the central nervous system that supports SGLT2i’s protective effects in treatment of cardiac remodeling

**DOI:** 10.1101/2025.10.02.674829

**Authors:** Miao Yuan, Hanxue Wu, Jiawei Wang, Zihan Qiu, Kexin Li, Jiaxi Xu, Dengfeng Gao

**Affiliations:** Department of Cardiology, Second Affiliated Hospital of Xi’an Jiaotong University, Xi’an 710004, Shaanxi, China; Department of Physiology and Pathophysiology, Xi’an Jiaotong University, Xi’an 710061, Shaanxi, China; Key Laboratory of Neural and Vascular Biology, Ministry of Education, Hebei Medical University, Shijiazhuang, Hebei, 050017, China

**Keywords:** Hypertension, Empagliflozin, Cardiac remodeling, Microglia

## Abstract

**Background:** Despite growing evidence of sodium-dependent glucose transporters 2 inhibitors (SGLT2i) improving heart failure, their mechanisms remain unclear. This study explored the effects of empagliflozin (EM) on sympathetic nerve activity and potential neuroimmune pathways.

**Methods:** Single-nucleus RNA sequencing (snRNA-seq) was performed in hypothalamic tissue of a DOCA-salt mouse model to screen for cellular changes. Validation of relevant mechanisms in DOCA-salt and high-salt diet (HSD) mouse models. Additionally, clinical data from 1,189 hospitalized heart failure patients were analyzed to evaluate the translational relevance of the experimental findings.

**Results:** EM treatment was associated with protective effects against cardiac remodeling in both DOCA-salt and HSD mice. Both models exhibited sympatho-excitation and neuronal hyperactivity in pre-autonomic brain regions, which appeared to be blunted by EM co-treatment. snRNA-seq analysis suggested that cellular interplay among vascular cells, microglia, and inhibitory neurons might be involved in these processes. Specifically, high-salt intake was linked to peripheral immune activation, including lymphocytosis and elevated plasma interferon-γ, which potentially mediated microglial state transitions. EM might counteract the disease-associated upregulation of protein ubiquitination. Clinically, SGLT2i use was significantly associated with reduced lymphocyte counts in specific patient subgroups.

**Conclusions:** Empagliflozin may attenuate cardiac remodeling caused by high sodium intake through neuroimmune effects by restoring microglial homeostasis and inhibiting sympathetic nerve activity.

## 1. Background

Cardiac remodeling refers to adaptive changes of the heart, in the perspective of structure and systolic function, following physiological or pathological stimuli(1). Recently, sodium-glucose co-transporter 2 inhibitors (SGLT2i) have shown beneficial effect in the treatment of heart failure(2, 3, 4). The mechanism of action of SGLT2i has been extensively studied, including its impact on cardiac metabolism, energy consumption, and direct effects on cardiac structure and function(5, 6).

Although multiple mechanisms of action of SGLT2i have been reported, the possibility of SGLT2i acting through the autonomic nervous system, especially the sympathetic nervous system, has not been fully dissected. The sympathetic nervous system plays a key role in the occurrence and development of cardiac diseases(7). It affects heart contractility and heart rate by releasing catecholamines such as epinephrine and norepinephrine. Beyond the heart, sympathetic nerves also innervate other organs that contribute to pathological development of cardiac remodeling, such as the kidney and spleen(8, 9, 10, 11, 12, 13). On one hand, elevation of renal sympathetic activity reduces renal blood flow, induces release of renin and aldosterone, thus increases water-sodium retention. On the other, elevation of sympathetic tone in the immune organs, leads to deployment of lymphocytes from the spleen(13), and enhances myeloid cell generation and leukocyte output in the bone marrow(14, 15). Renin-angiotensin-aldosterone system (RAAS) has been shown to induce pro-inflammatory transformation of leukocytes and aggregation of T lymphocytes(16), to exacerbate inflammatory responses. Within the brain, the paraventricular nucleus of hypothalamus (PVN), a pre-autonomic center, has high capillary density, thin vessel diameter, and complex vascular topology(11, 17, 18, 19, 20, 21), providing a possible interface for immune cells and cytokines to further boost sympathetic outflow. Accordingly, integrated effect of blunted sympatho-excitation to these organs could decrease the cardiac afterload, support the reduction of cardiac energy consumption, and attenuate inflammation, therefore blunting pathological development of cardiac remodeling. Interestingly, SGLT2i increases urinary sodium excretion, reduces renal blood flow, stimulates short-term renin secretion, but long-term impact on renin and aldosterone is not observed(22, 23). Clinically, SGLT2i is often applied with diuretics or RAAS inhibitors(23). Investigating how SGLT2i alone affect central regulation of sympathetic tone would provide novel insight regarding its mechanism of action in cardiac remodeling, and a new perspective on its clinical application.

This study used deoxycorticosterone acetate (DOCA)-salt and high-salt (8%) diet (HSD) mouse models, along with blood samples from heart failure patients, to examine the effects of SGLT2i on the peripheral immune and central nervous systems. We aimed to study how SGLT2i regulates signaling from leukocytes to pre-autonomic neurons, suggesting a potential neurogenic mechanism for improving cardiac remodeling.

## 2. Results

### 2.1 Empagliflozin attenuates pathological cardiac remodeling induced by salt overload

In this study, the therapeutic effect of empagliflozin (EM) was studied in both DOCA-salt model and 8% high-salt diet model (HSD). DOCA-salt model (3-week) was utilized for its character that induces salt-sensitive type of hypertension and does not elevate circulating renin and angiotensin levels(24, 25, 26, 27), while HSD (8-week) was used because it increases salt-intake but cannot induce hypertension in C57BL6/J mice(28, 29, 30). To ensure that the observed benefits of EM were not merely secondary to its modest BP-lowering effect, hydralazine was administered to a separate group (DOCA+Hyd) to serve as a normotensive control for the DOCA-salt model. Firstly, there was no significant difference in body weight among the five groups of mice received treatment of DOCA-salt, DOCA+Hyd, DOCA+EM, sham, and sham+EM, respectively (Supplementary Figure1 A). Similarly, no significant difference in body weight was observed among the groups with HSD or normal salt diet (NSD) treatment (Supplementary Figure1 B). EM co-treatment did not affect drinking water consumption in DOCA-salt-treated mice but slightly increase water intake in mice with HSD (Supplementary Figure1 C-F). Baseline blood pressure (BP) was comparable across all groups before treatment (Supplementary Figure 1 G-J). In DOCA-salt-treated mice, EM co-treatment did not alter either systolic or diastolic BP; thus, mice in the DOCA+EM group still exhibited hypertension. In contrast to EM, hydralazine significantly lowered BP in DOCA-salt-treated mice (Supplementary Figure 1 K, L). HSD did not induce hypertension, and EM co-treatment had no effect on BP in this model (Supplementary Figure 1 M, N).

Cardiac remodeling was evaluated in the DOCA-salt model. Firstly, we observed increased ratios of heart weight to body weight and heart weight to tibia length in DOCA-salt mice (Figure 1A, B), indicating the development of cardiac hypertrophy. Histological staining showed that DOCA-salt treatment induced cardiomyocyte hypertrophy and increased both interstitial and perivascular fibrosis in the left ventricle (LV) (Supplementary Figure 2 A-C). WGA staining showed that EM could passivate DOCA-salt induced cardiomyocyte hypertrophy (Figure 1 C, F). These changes were accompanied by reduced left ventricular ejection fraction (LVEF), increased LV mass (Figure 1 C-E), and significant Collagen III deposition (Figure 1 G). EM co-treatment significantly inhibited LV hypertrophy, reduced fibrosis and Collagen III deposition, and preserved cardiac systolic function. Notably, hydralazine failed to show similar protective effects against pathological cardiac remodeling.

Similar results were observed in the HSD model. EM treatment improved HSD-induced increases in heart weight ratios (Figure 1 H, I), cardiomyocyte hypertrophy, and tissue fibrosis (Figure 1J, M, Supplementary Figure 2 D-F). Furthermore, HSD did not lead to a significant decrease in LVEF, and EM could improve the increase in left ventricular mass caused by HSD (Figure 1 J-L), while reducing collagen deposition. (Figure 1 N). Collectively, these results demonstrate that EM protects against pathological cardiac remodeling induced by DOCA-salt or HSD treatment. These findings suggest that EM improves cardiac remodeling in both salt-sensitive (DOCA-salt) and salt-insensitive (HSD) models. Given that high salt intake itself can directly impair cardiac structure, the promotion of sodium excretion by EM may contribute to its protective role, independent of its blood-pressure-lowering effects.

Since EM increases urinary sodium excretion, we measured plasma sodium levels and osmolality to determine if its cardiac protection was mediated by lowering these parameters. No significant differences were observed in plasma sodium or osmolality across groups in the DOCA-salt model and the HSD model (Supplementary Figure 3 A-D). However, DOCA-salt treatment significantly reduced hematocrit (HCT) levels, indicating increased plasma volume (Supplementary Figure 3 E). Notably, EM co-treatment, but not hydralazine, restored HCT to baseline levels. Unlike the DOCA-salt model, HSD did not induce changes in HCT (Supplementary Figure 3 F).

### 2.2 Empagliflozin blunts sympathetic hyperactivity

To evaluate systemic sympathetic activity, we first measured urinary norepinephrine (NE) levels normalized by creatinine (Cr). In both salt-sensitive (DOCA-salt) and salt-insensitive (HSD) models, excessive sodium intake significantly increased urinary NE levels (Figure 2 A, B). Notably, this systemic activation was significantly reduced by EM co-treatment, whereas hydralazine showed no inhibitory effect, even though it lowered blood pressure.

To further investigate whether these systemic changes were reflected in the heart, we assessed cardiac sympathetic innervation by measuring tyrosine hydroxylase (TH) expression in representative groups (Sham, DOCA-salt, and DOCA+EM). We focused on these groups to specifically evaluate the blood-pressure-independent effects of EM on local sympathetic tone. DOCA-salt treatment markedly increased the proportion of TH-positive cardiomyocytes in the left ventricle (LV), a change that was significantly blunted by EM co-treatment (Figure 2 C, D). Similarly, in the HSD model, EM effectively attenuated the increase in LV TH expression induced by high sodium intake (Figure 2 E, F).

Furthermore, we examined the expression of c-Fos, a marker of neuronal activation, within the paraventricular nucleus (PVN) and the rostral ventrolateral medulla (RVLM), which are critical pre-autonomic centers. Consistent with the peripheral sympathetic activation, high sodium intake induced a marked increase in c-Fos-positive neurons in both the DOCA-salt and HSD models. Notably, EM co-treatment effectively blunted this neuronal hyperactivity in both models (Figure 3). Given that EM suppressed neuronal over-excitation even in the normotensive HSD model, these findings suggest that the neuroprotective effect of EM on central autonomic centers is likely independent of systemic blood pressure regulation.

### 2.3 snRNA-seq Reveals Vascular-Immune Alterations in the PVN

To explore the cellular transcriptomes of the PVN under EM treatment, we performed single-nucleus RNA sequencing (snRNA-seq) on tissues from Sham, DOCA-salt, and DOCA+EM groups (Figure 4A). After quality control and clustering, we identified 37, 828 nuclei, categorized into nine major cell types, including neurons (excitatory, inhibitory, and mixed), glia (immature oligodendrocytes, mature oligodendrocyte, astrocytes, and microglia), ependymal, and vascular cells (endothelial cells & pericytes) (Figure 4 B, C). Markers such as *Sim1* confirmed the precise capture of the PVN area (Supplementary Figure 4).

To identify potential cellular interactions associated with EM treatment, we performed CellChat analysis. In the DOCA-salt group, we observed the emergence of predicted signaling from endothelial cells and pericytes to microglia, an interaction that was absent in the Sham group. Furthermore, while the crosstalk between microglia and inhibitory neurons was present in both Sham and DOCA-salt groups, this specific interaction was effectively abolished by EM co-treatment (Figure 4 D). These results suggest that EM may exert its protective effects by selectively interrupting pathological communication between reactive microglia and pre-autonomic inhibitory neurons.

Differentially expressed gene (DEG) analysis (Figure 4 E) and Gene Ontology (GO) enrichment (Figure 4 F) further revealed that transcriptional changes in endothelial cells and pericytes induced by both DOCA-salt and EM were significantly linked to cytokine signaling and immune cell adhesion. These results suggest that DOCA-salt may induce an immune response in PVN vessels, potentially increasing the recruitment of circulating leukocytes—an effect that was attenuated by EM. Since the PVN has a dense capillary network that serves as an interface for systemic-brain communication(11, 17, 18, 19, 20, 21). It has been shown that cytokines from leukocytes, such as interferon, has direct impact on microglia(31).We next analyzed the microglial population to evaluate their response to these peripheral signals.

### 2.4 EM suppresses microglial reactive transformation and peripheral inflammation

To further investigate microglial states, we sub-clustered the microglia into four groups based on their transcriptional profiles: homeostatic (HOM), disease-associated (DAM), interferon-activated (IFN), and oligodendrocyte related microglia (ORM) microglia (Figure 5A-C). This fine-grained analysis allowed us to pinpoint the specific microglial responses to salt overload and EM treatment.

Within the HOM and DAM clusters, DOCA-salt treatment was associated with the enrichment of genes involved in microglial activation and phagocytosis. Conversely, EM co-treatment modulated genes linked to cell adhesion and oxidative phosphorylation, suggesting a metabolic and functional shift in microglial states (Figure 5D-G). Furthermore, the IFN-activated microglia cluster, which is highly sensitive to peripheral cytokines, exhibited distinct transcriptional profiles. Specifically, DOCA-salt treatment upregulated genes involved in efferocytosis, whereas EM treatment predominantly modulated phagocytic pathways (Figure 5H, I). Notably, several IFN-responsive genes, such as Txndc9, Ppia, and Adam17, showed reciprocal expression patterns between the DOCA-salt and EM groups, identifying them as potential molecular targets for EM’s neuroprotective effects (Figure 5J).

To validate these transcriptomic shifts at the protein level, we performed immunofluorescence staining in the paraventricular nucleus (PVN). Consistent with our snRNA-seq findings, EM significantly attenuated the DOCA-salt-induced accumulation of Iba1 /CD68 reactive microglia (Figure 6A, B). Similarly, EM blunted the microglial reactive transformation triggered in the HSD model (Figure 6 C). Finally, we assessed the peripheral immune response to determine if these central changes were mirrored systemically. Both DOCA-salt and HSD treatments significantly increased circulating lymphocyte counts and plasma IFN-γ levels. EM co-treatment effectively normalized these peripheral inflammatory markers in both models (Figure 6 D-G). Additionally, EM mitigated the increase in neutrophil counts caused by high sodium intake (Supplementary Figure 3 G, H).

The consistency of these immune profiles across both DOCA-salt and HSD models reinforces the notion that excessive sodium intake, regardless of the specific hypertensive model, triggers systemic and central immune activation. Collectively, these results demonstrate that EM treatment attenuates both peripheral inflammation and the subsequent reactive transformation of microglia in the PVN, providing a potential mechanism for its neuroprotective and cardioprotective effects.

### 2.5 EM blunts protein ubiquitination and preserves ACE2 activity in PVN neurons

Next, we analyzed snRNA-seq data from PVN neurons. In both excitatory and inhibitory neurons, DOCA-salt treatment upregulated the protein ubiquitination pathway, an effect that was significantly reversed by EM co-treatment (Figure 7 A, B). These findings suggest that EM may "stabilize" PVN neuro-endocrine function by modulating protein degradation.

Previous studies have shown that ACE2 ubiquitination impairs its enzymatic activity, thereby potentially compromising the GABAergic inhibitory tone in the PVN. To investigate this, we measured ACE2 activity in the hypothalamus. DOCA-salt and HSD treatments markedly reduced ACE2 enzymatic activity, whereas EM co-treatment effectively preserved or restored it (Figure 7 C, D). Furthermore, we examined NEDD4L, the E3 ligase responsible for ACE2 ubiquitination. Both DOCA-salt and HSD models exhibited increased NEDD4L expression, which was significantly blunted by EM co-treatment (Figure 7 E-H).

To identify the specific neuronal types responding to EM, we analyzed PVN-specific markers. DOCA-salt treatment upregulated the expression of *Oxt, Avp, Tac1, Trh,* and *Sst*, while EM co-treatment downregulated these markers (Figure 7 I), suggesting the involvement of neuro-endocrine and neurosecretory neurons. Consistently, the modeling-induced elevation of plasma copeptin, a reliable surrogate for arginine vasopressin (AVP), was markedly blunted by EM (Figure 7 J, K). Collectively, these results suggest that the protective effects of EM are associated with the suppression of the NEDD4L/ACE2 ubiquitination pathway and the subsequent stabilization of neuro-endocrine activity.

### 2.6 EM is associated with altered hematocrit and lymphocyte counts in heart failure patients

To evaluate the clinical relevance of our experimental findings, we conducted a retrospective cross-sectional study involving patients hospitalized with heart failure at the Second Affiliated Hospital of Xi’an Jiaotong University between 2018 and 2023. Of the 1,864 screened patients 1,189 met the inclusion criteria (Non-treated: n = 715; SGLT2i-treated: n = 474). After Propensity Score Matching (PSM) to balance baseline clinical characteristics (Table 1, Figure 8), systemic inflammatory and volume-related markers were compared between the matched cohorts. Post-PSM analysis revealed that the SGLT2i-treated group had significantly higher hematocrit (HCT) levels (*P* < 0.001) and lower counts of lymphocytes and monocytes (*P* < 0.001 and *P* = 0.02, respectively) compared to the non-treated group (Table 2). To account for remaining baseline differences (e.g., alcohol consumption, NT-proBNP levels, and diuretic use), we performed stratified analyses.Regardless of the clinical strata, HCT levels remained consistently higher in the SGLT2i-treated group. Notably, significantly lower lymphocyte counts were observed in specific SGLT2i-treated subgroups, including patients who did not consume alcohol, those with high NT-proBNP levels (> 2550 pg/mL), and those not receiving diuretics (*P* < 0.001). Additionally, SGLT2i was associated with lower neutrophil counts in patients concurrently treated with diuretics (*P* = 0.004) (Figure 8).In summary, our clinical data indicate that SGLT2i treatment is associated with a specific hematological and immune profile—characterized by increased HCT and reduced circulating inflammatory cells—which mirrors our observations in mouse models and suggests its potential role in modulating systemic inflammation and volume status in heart failure patients.

**Table 1.**
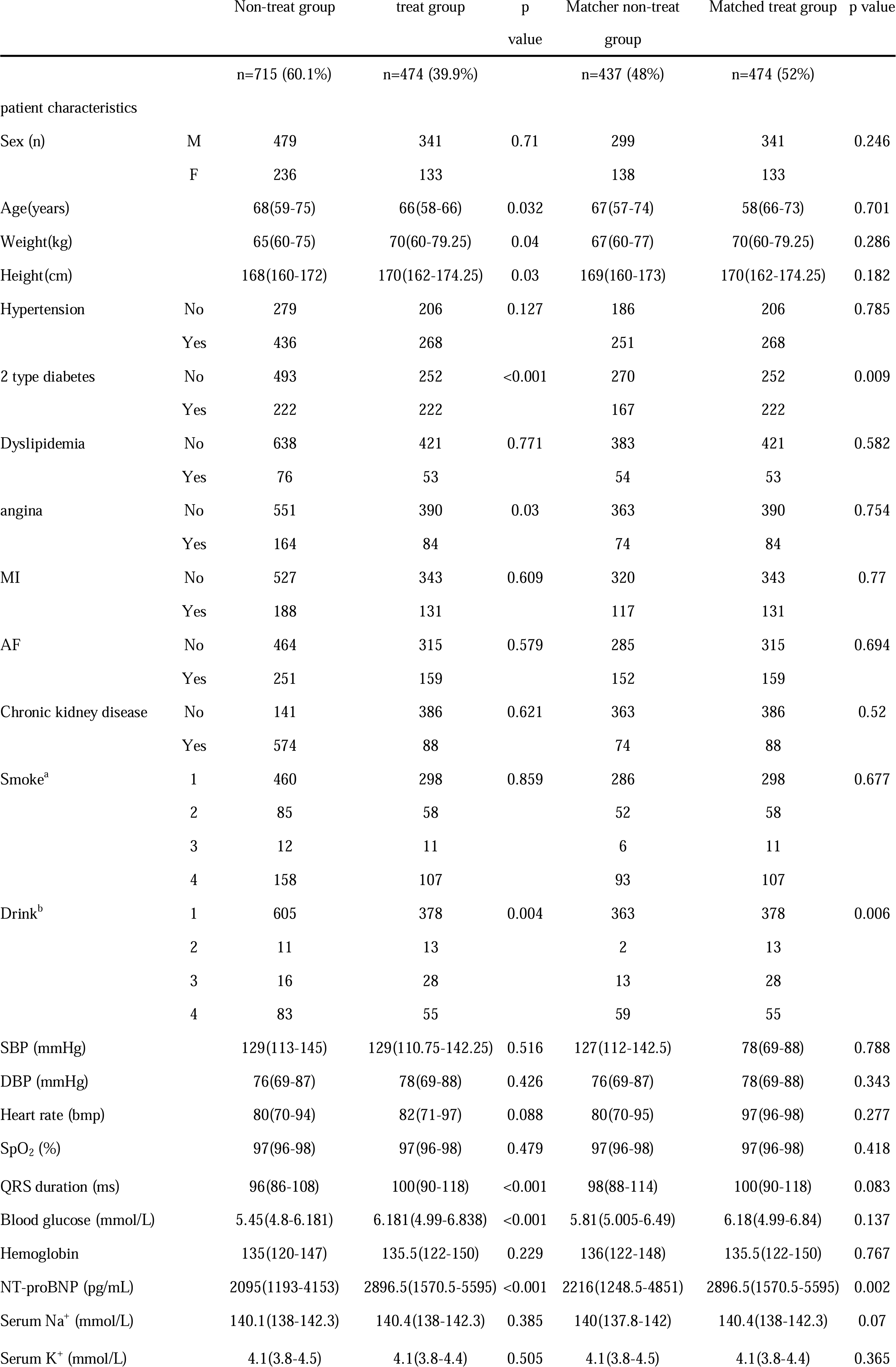
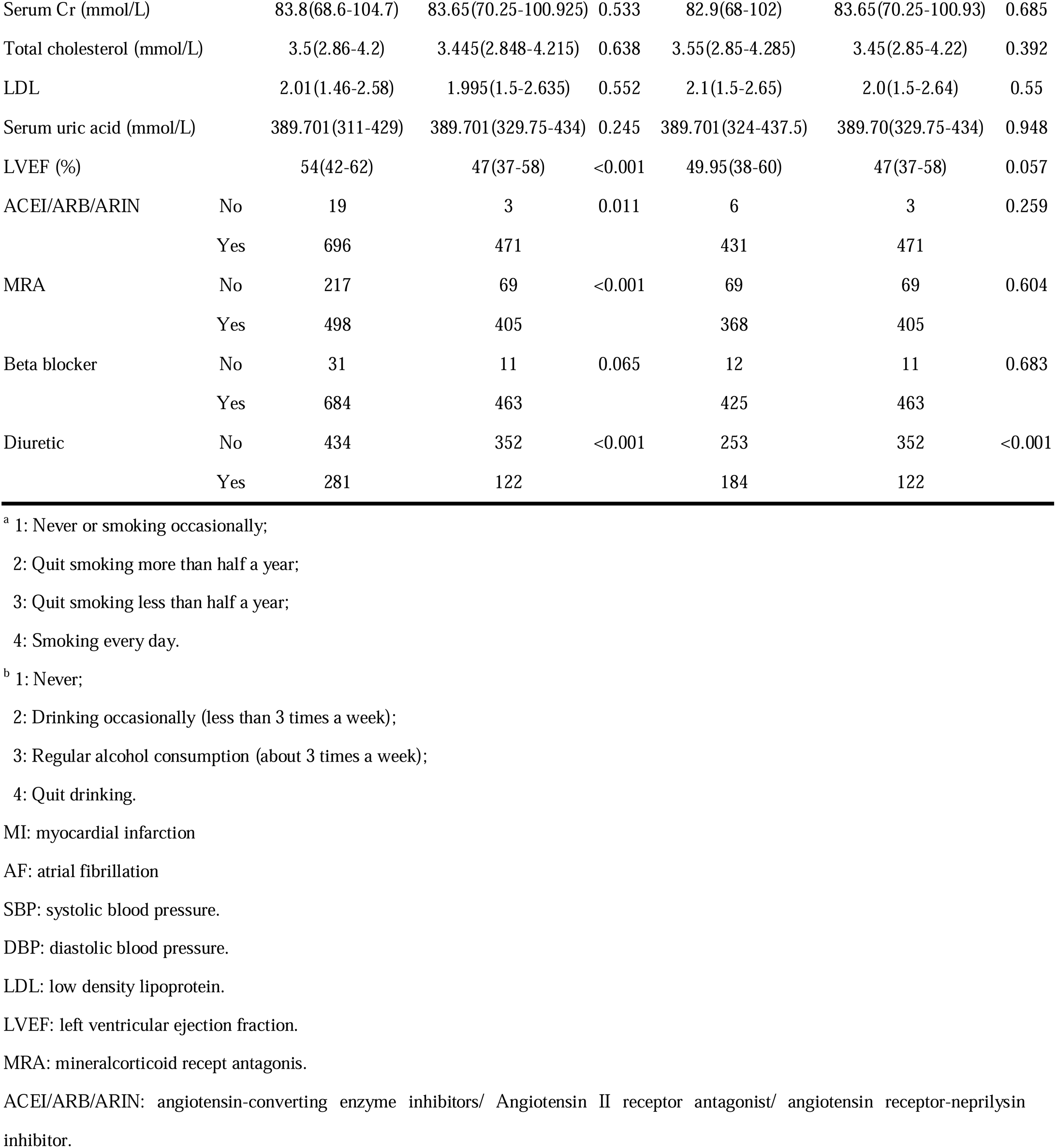
Baseline comparison of the treated group and non-treated group before and after propensity score matching.

**Table 2.** Comparison of HCT and peripheral immune cell counts between the treated group and non-treated group after propensity matching score.

|  | Matched non-treat group | Matched treat group | *P* value |
| --- | --- | --- | --- |
| n=911 | n=437 (48%) | n=474 (52%) |  |
| HCT | 39.9(37-43.4) | 43(39.63-46.1) | <0.001 |
| Neutrophil count | 4.29(3.34-5.58) | 4.18(3.19-5.31) | 0.22 |
| Monocyte count | 0.52(0.4-0.68) | 0.49(0.39-0.63) | 0.02 |
| Lymphocyte count | 1.52(1.19-1.91) | 1.39(1-1.77) | <0.001 |

## 3. Discussion

Emerging evidence highlights cardiovascular protective effects of SGLT2i, potentially linked to sympathetic inhibition (32), yet their impact on autonomic brain centers remains largely unexplored. In this study, using both DOCA-salt and 8%-HSD models, we demonstrated that EM protects against cardiac remodeling, at least in part, by suppressing central sympathetic outflow. Our findings suggest that EM exerts its protective role by attenuating peripheral cytokine-mediated microglial activation and preventing the over-excitation of neuro-endocrine and neurosecretory neurons (Figure 9). The clinical and experimental data collectively point to an important role for the immune system in EM’s protective effects. NT-proBNP is an important diagnostic biomarker for of acute heart failure with higher mortality risk. Inflammation appears to be associated with natriuretic peptide (NP) release(33). In heart failure, BNP levels closely correlate with the extent of left ventricular sympathetic overactivity (34, 35).

We found that in patients with high NT-proBNP levels—who are typically at higher clinical risk—EM treatment significantly reduced circulating lymphocyte counts. This clinical observation mirrors our findings in mice, where EM reduced lymphocyte and neutrophil counts in both salt-related models (Supplementary Figure 3 G, H). While neutrophils may contribute to cardiac pathology, their recruitment to the brain is limited under non-infectious conditions. Therefore, we hypothesize that lymphocytes and their associated cytokines serve as the primary mediators of systemic-to-brain communication in this context. We previously showed that DOCA-salt increases T cell ratios in peripheral blood, and targeting the brain to ablate sympatho-excitation reduces these ratios (36).

EM has been shown to reduce inflammation in diabetic patients (37, 38, 39, 40). In the current study, EM treatment lowered both lymphocyte counts and plasma levels of IFN-γ. Given that IFN-γ is closely linked to hypertension progression (16, 41, 42), and can directly act on CNS microglia (43, 44, 45) to promote neuroinflammation, EM likely blunts sympathetic outflow by mitigating this peripheral-to-central inflammatory cascade. Reactive microglia in the PVN have been demonstrated to promote excitation of PVN neurons and increase sympathetic outflow (11, 17, 18, 19, 20, 21).

In addition to immune modulation, EM significantly impacts water-sodium homeostasis and neuronal protein stability. EM increased HCT in both DOCA-salt mice and heart failure patients, a change often associated with reduced blood volume due to SGLT2i-induced diuresis (46, 47, 48, 49, 50, 51). However, EM also improved outcomes in patients already receiving diuretics, suggesting additional effects beyond simple fluid clearance. While blood sodium and osmolality remained stable due to robust systemic regulation (Supplementary Figure 3 A-D), we observed significant changes in neuro-endocrine markers. Vasopressin, secreted by PVN neurosecretory AVP neurons, is essential for the development of DOCA-salt hypertension (24). We found that EM restored plasma co-peptin levels (a surrogate for AVP) to baseline in both models, further supporting the involvement of the CNS. This stabilization of AVP neurons may be linked to the ACE2/NEDD4L pathway. Our snRNA-seq data revealed that EM suppressed the protein ubiquitination pathway in PVN neurons (Figure 7 A, B). Specifically, EM blunted the expression of NEDD4L (an E3 ligase) and preserved ACE2 enzymatic activity, which is known to be impaired by NEDD4L-mediated ubiquitination during hypertension. By facilitating sodium excretion and potentially stabilizing the NEDD4L/ACE2 axis, EM reduces the over-excitation of neuro-endocrine neurons, thereby providing multi-level protection against pathological remodeling.

## 4. Limitations and Conclusion

This study has several limitations that should be acknowledged. The clinical portion utilized a single-center retrospective design, which may introduce selection bias or unmeasured confounding factors despite the use of propensity score matching. Furthermore, the reliance on retrospective medical records might lead to the exclusion of severe cases or those lost to follow-up, potentially underestimating certain adverse outcomes. Nevertheless, the consistency between our clinical observations and experimental findings supports the validity of our conclusions. In summary, our study provides novel evidence that EM suppresses pathological cardiac remodeling by modulating the peripheral-central immune axis and stabilizing neuro-endocrine activity in the PVN. These effects appear to be independent of its conventional anti-hypertensive actions, offering a broader therapeutic perspective for the use of SGLT2i in cardiovascular diseases.

## 5. Methods

### 5.1 Antibodies

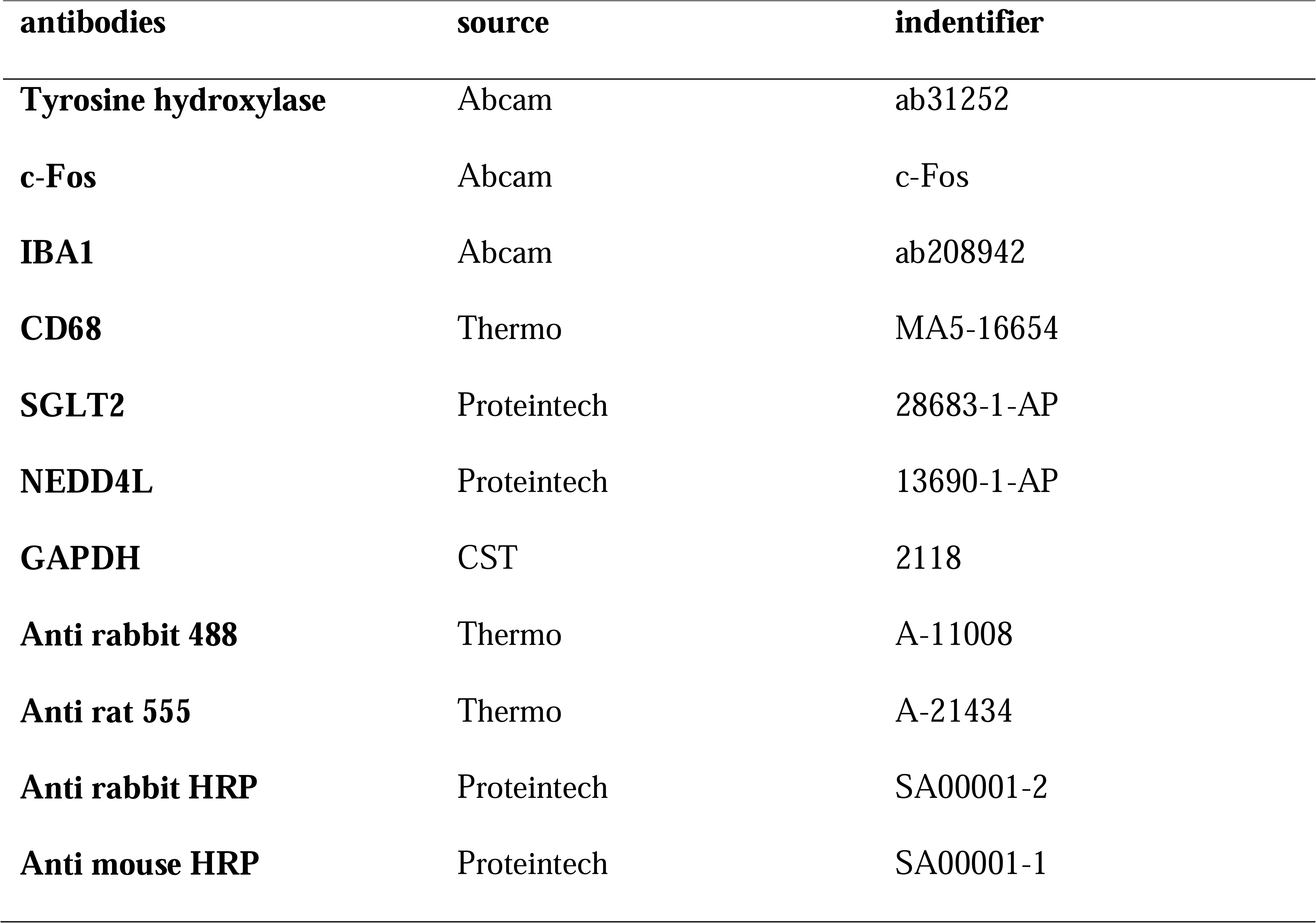

| antibodies | source | identifier |
| --- | --- | --- |
| **Tyrosine hydroxylase** | Abcam | ab31252 |
| **c-Fos** | Abcam | c-Fos |
| **IBA1** | Abcam | ab208942 |
| **CD68** | Thermo | MA5-16654 |
| **SGLT2** | Proteintech | 28683-1-AP |
| **NEDD4L** | Proteintech | 13690-1-AP |
| **GAPDH** | CST | 2118 |
| **Anti rabbit 488** | Thermo | A-11008 |
| **Anti rat 555** | Thermo | A-21434 |
| **Anti rabbit HRP** | Proteintech | SA00001-2 |
| **Anti mouse HRP** | Proteintech | SA00001-1 |

### 5.2 Stored data

| Database | Number |
| --- | --- |
| **GEO dataset** | GSE313832 |
| **SRA dataset** | SUB15305597 |

### 5.3 Software and algorithms

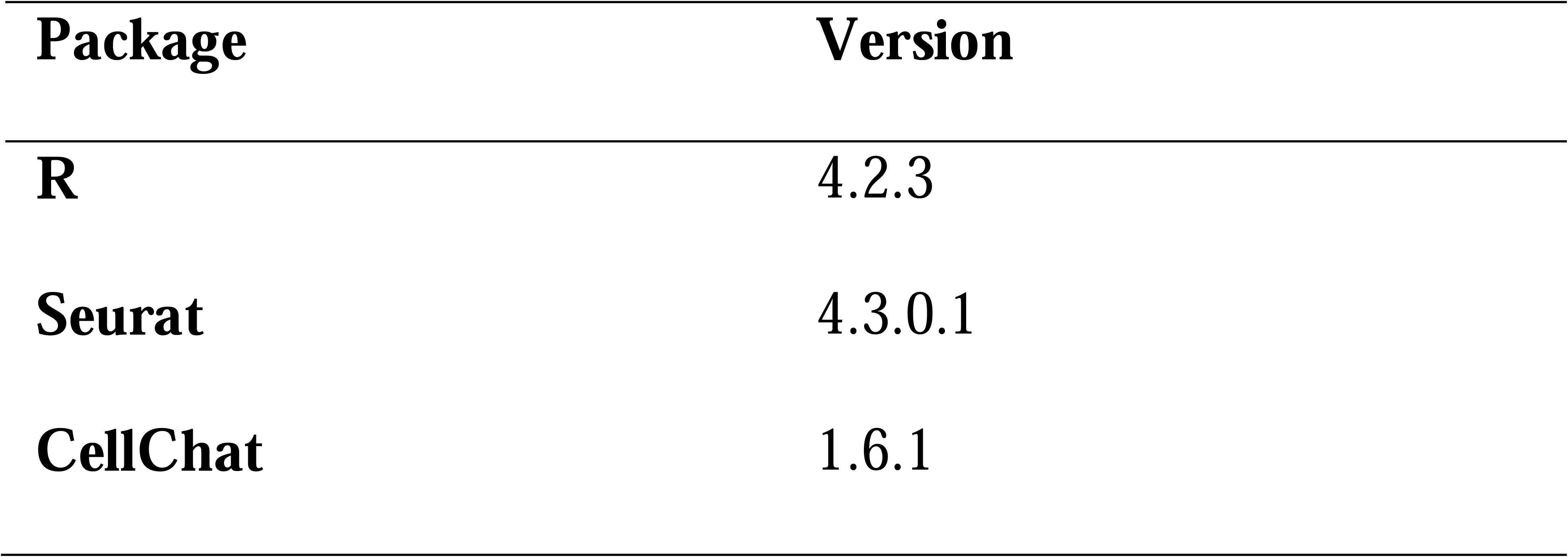

| Package | Version |
| --- | --- |
| R | 4.2.3 |
| Seurat | 4.3.0.1 |
| CellChat | 1.6.1 |

### 5.4 Experimental Model and Subject Details

This study used male C57BL/6J mice of SPF level at 8 weeks old. They were purchased from Beijing Vital River Laboratory Animal Technology Co., Ltd. (Order No. 44829700009980). The animals were raised in the Experimental Animal Center of Xi’an Jiaotong University under SPF barrier conditions. The animal feed was purchased from Jiangsu Xietong Pharmaceutical Bio-engineering Co., Ltd. (regular feed: SWS9102, 8% NaCl feed: XTNacl-8C). Mice had free access to water and food. All procedures were approved by the IACUC of Xi’an Jiaotong University (XJTU2021-203), in agreement with the Guide for the Care and Use of Laboratory Animals. Every effort was made to minimize the number of animals used and their suffering. Institutional Animal Care and Use Committees in accordance with the ‘Principles of Laboratory Animal Care by the National Society for Medical research and the Guide for the Care and Use of Laboratory Animals’ (National Institutes of Health Publication No. 86-23, revised 1996).

### 5.5 DOCA-salt Model and High Salt Diet Model

All mice were adaptively fed for 1 week. The DOCA group, DOCA+EM group, DOCA+Hyd group, Sham+EM group and Sham group all underwent right kidney removal. The DOCA group, DOCA+EM group and DOCA+Hyd group underwent Deoxycorticosterone acetate (DOCA) silicone implantation. Anesthesia was induced with isoflurane (3% induction; 1-2% maintenance) using an oxygen flow of 0.5 L/min, and the mice were placed on a heating pad to maintain body temperature at around 37.5. The mouse was placed in a left lateral position on the operating table, and the skin in the posterior peritoneal area was incised to expose the renal hilum of the right kidney. The blood vessels were ligated with surgical suture at the renal hilum, followed by removal of the right kidney. After surgery, subcutaneous injection of 0.9% saline solution (1mL) was given via backpack skin. One week after recovery, either DOCA, DOCA+EM or DOCA+Hyd groups received subcutaneous implantation of DOCA silicone sheet (containing 1 mg of DOCA per gram body weight). Mice implanted with DOCA silicone sheets had their drinking water replaced with 1% NaCl solution.

HSD, HSD + EM and normal sodium diet (NSD) groups were fed adaptively for one week. HSD, HSD + EM were fed with a diet containing 8% sodium for 12 weeks. The NSD group was fed with a normal diet (sodium content was 0.45%) for 12 weeks. All mice were given free access to food and water.

All mice treated with EM were administered by daily gavage at a dose of 10mg/kg/day. The dosages of the EM were selected based on the literature. All tissue preparations were performed after euthanasia. Using a precision vaporizer with induction chamber and waste gas scavenger, (indicate the gas anesthetic) will be administered slowly up to [indicate: > 4.5 % (for Isoflurane)] in oxygen and continued until respiratory arrest occurs for > 60 seconds. The chamber is flushed with oxygen only, the animal is removed and rapid removal of (indicate tissues / organs) is performed to assure euthanasia.

### 5.6 Single-cell RNA-sequencing

5 mice were sacrificed with a lethal dose of isoflurane, and the excess blood was washed off in cold PBS, and the PVN region of each mouse was obtained according to the mouse brain anatomy map. The tissue nucleus was extracted by SHBIO tissue sample nucleus isolation kit (Cat# 52009-10).

Single-cell RNA-Seq libraries were prepared using SeekOne^®^ Digital Droplet (SeekOne^®^ DD) Single Cell 3’ library preparation kit (SeekGene Catalog No. K00202-0201). The indexed sequencing libraries were cleanup with SPRI beads, quantified by quantitative PCR (KAPA Biosystems KK4824) and then sequenced on Illumina NovaSeq 6000 with PE150 read length or DNBSEQ-T7 platform with PE150 read length.

We used the SeekSoul Tools pipeline to process the cleaned reads and generated the transcript expression matrix. Quality control, clustering, and downstream analysis were performed using Seurat v. 4.3.0.1 in R version 4.2.3 (2023-03-15). Briefly, according to gene, UMI, and mitochondrial gene counts (genes > 100 and < 8000, mitochondria < 5%).

For the clusters, data was normalized and scaled using the NormalizeData function in Seurat with a factor of 10,000. The top 3000 variable genes were selected using the FindVariableFeature function in Seurat, and the data was scaled using the ScaleData function. Cells were clustered using the FindCluster function with 20 principal components and a resolution of 1.0. The cells were then plotted in a two-dimensional space using the ‘Unified Manifold Approximation and Projections’ (UMAP) technique, and the identities of the cell clusters were determined by their overlap with known marker genes in the literature. 35 clusters were identified as 9 cell types: Mature oligodendrocyte (5,854), Excitatory neuro (11,225), Inhibitory neuro (11,390), Mix neuro (4,620), Astrocyte (749), Microglial cell (1444), Immature oligodendrocyte (1,264), Endothelial cell & Pericyte (176), Ependymal cell (106). In order to better describe the neuron population, excitatory neurons, inhibitory neurons, and mixed neurons were further separated and re-clustered. Cells were clustered using the FindCluster function with 20 principal components and a resolution of 0.1.

We performed cell-cell communication analysis using CellChat v1.4.0, which is based on the curated ligand-receptor interaction database (CellChatDB). To perform the analysis, we provided normalized gene expression matrices for PVN tissues as input to CellChat. The computeCommunProb function was used to calculate the total numbers of interactions and interaction strengths, while the computeCommunProbPathway function was used to calculate the communication probabilities for each cell signaling pathway.

### 5.7 Echocardiography

Transthoracic echocardiography was performed 21 days after DOCA silicone block implantation. Echocardiography was performed under anesthesia with isoflurane (0.25 to 0.50%) supplemented with 100% O2 using a 30MHz probe on Vevo^®^LAZR-X 3100. Left ventricular mass (LVM) and left ventricular ejection fraction (LVEF) were analyzed from M-mode images.

### 5.8 Staining of the Heart

After the mice were directly sacrificed, the heart was removed, and the blood washed off the surface in cold PBS. The heart was placed in 4% PFA for 48 hours and then paraffin-embedded sections (10 μm) were performed. The H&E staining kit (Servicebio Cat#: G1076-500ML) and MASSON staining kit (Servicebio Cat#: G1006-20M) were used.

### 5.9 Immunocytochemistry

After antigen repair with sodium citrate, the paraffin sections of the heart were washed twice with PBS, then 10min was incubated with 0.25% TritonX100 at room temperature, and any non-specific binding was blocked by 10% NGS at room temperature for 1 hour and incubated overnight with TH primary antibody (1:1000, 4 □). After washing with PBS for three times, the rabbits-488 were incubated at room temperature for 1 hour and then washed again with PBS. Finally, anti-staining was performed with DAPI (1-VOL1000-THERMOTIMO-Fisher Science).

For brain tissue, mice were fixed by cardiac perfusion with cold 4% PFA in PBS. After the brain tissue was fixed by 4% PFA and embedded with OCT (SAKURA-4583), the brain map skin of the control mice was sliced (25 μm) and specific parts were taken for immunofluorescence staining. Wash with PBST twice, then incubate 10 min with 0.5% Triton-X100 at room temperature, block any non-specific binding with 10% NGS at room temperature for 1 hour, dilute the primary antibody in proportion and incubate overnight with 4. After washing with PBS for three times, the corresponding secondary antibody was used to incubate at room temperature for 1 hour, and then cleaned again with PBS. The cell nucleus was stained with DAPI (Thermo Fisher Science Cat#: 62247). The main antibodies used in this study are summarized in in. Finally, the fluorescence microscope (Olympus BX43) was used to capture the image, and the ImageJ software was used for image analysis.

### 5.10 Norepinephrine Measurement

Urine samples were collected by bladder massage at the end of the experimental protocol and noradrenaline concentrations were determined using a mouse noradrenaline ELISA kit (Elabscience, Cat#: E-EL-0047) according to the manufacturer’s instructions. Noradrenaline levels are normalized by the corresponding creatinine concentration (Elabscience, Cat#: E-BC-K188-M). Normalized noradrenaline concentrations in urine are expressed as ng/ng creatinine.

### 5.11 Peripheral Blood Cell Counts and Plasma Sodium Detection

After the experimental protocol was completed, the mice were anesthetized with isoflurane and blood was taken through the eyeball. Half of blood was anticoagulation by EDTA anticoagulation. Then the blood was routinely examined by a fully automated three-point population blood cell analyzer (Mindray BC-2800). The remaining blood was anticoagulated by EDTA and centrifuged for 4 □. Following the manufacturer’s instructions to use blood sodium detection kit (Elabscience, Cat#: E-BC-K207-M), MCP-1 test kit (Elabscience, Cat#: E-EL-M3001), co-peptin test kit (JINGMEI BIOTECHNOLOGY, Cat#: JM-12422M1), and IFN-γ test kit (Elabscience, Cat#: E-EL-M0048) to determine the corresponding indicators. The osmotic pressure was detected by an osmometer (Fiske Micro-Osmometer Model 210).

### 5.12 ACE2 Enzyme Activity Detection

Mice were sacrificed with a lethal dose of isoflurane, and the paraventricular nucleus region of the inferior thalamus was obtained for each mouse according to the mouse brain anatomy map. ACE2 enzyme activity was determined using the ACE2 enzyme activity detection kit (Beyotime, Cat#: P0319S) according to the manufacturer’s instructions.

### 5.13 Western Blotting

The tissue samples were placed in 8M urea solution (4.8g urea, 10% SDS 5mL 0.5mL 1M Tris PH 7.4 0.5mL, 0.5M EDTA 0.1mL plus H_2_O to 10mL) supplemented with a mixture of protease inhibitors (Roche, Cat#: 11697498001) and repeatedly pumped with a 1ml needle. After quantifying the protein concentration by BCA protein assay (Thermo Scientific, Cat#: A55860), the protein sample was mixed with 4 × SDS sample buffers (Thermo Scientific, Cat#: NP0007) and boiled at 65 °C for 10 minutes. 30 μg protein samples were separated on 4-12% MOPS gel (ACE, Cat#: F15412LGel) and transferred to PVDF membrane (Millipore). Seal the imprint in 5% skim milk for 1 hour. On the second day, the protein was detected with specific antibody (diluted by 1purl 1000) and secondary antibody conjugated with HRP (diluted by 1purl 2000). Aim at the rabbit secondary antibody conjugated with NEDD4L and HRP conjugated with GAPDH.

The ECL luminescent solution (MISHUSHENGWU, Cat#: MI00607B) was dripped evenly on the PVDF membrane (Millipore, Cat#: IPVH 00010) and the imaging was carried out by the WB experimental chemiluminescence instrument (Bio-rad, Cat#:0087170). The signal strength was quantified by ImageJ.

### 5.14 Clinical Study Design

This study is a retrospective single-center clinical study. All participants provided written informed consent before participating in the study. The study was approved by the Institutional Review Board at the Second Affiliated Hospital of Xi’an Jiaotong University (IRB#2024-150). All methods were performed in accordance with the relevant guidelines and regulations.

#### 5.14.1 Study Participants

We retrospectively reviewed patients diagnosed with heart failure who were hospitalized at the Second Affiliated Hospital of Xi’an Jiaotong University between January 2018 and January 2023. The inclusion criteria were: (1) aged >18 years with signed informed consent; and (2) a diagnosis of heart failure within 3 months prior to enrollment. Patients were excluded based on the following criteria: (1) history of cerebrovascular accident; (2) fever of various etiologies or severe infection; (3) moderate-to-severe anemia; (4) uncontrolled hypertension (systolic blood pressure >160 mmHg); (5) renal failure; (6) hyperthyroidism or other significant endocrine diseases; and (7) SGLT2i treatment duration of less than one month.

A total of 1,864 patients were initially screened, of whom 1,189 were enrolled in the final study population, including 474 patients in the SGLT2i-treated group and 715 in the non-treated group. To balance the baseline clinical characteristics between the two cohorts, propensity score matching (PSM) was performed as detailed in the Statistical Analysis section. Clinical profiles and hematological outcomes were compared both before and after matching. The primary endpoints of this study were hematocrit (HCT), and peripheral inflammatory cell counts, specifically neutrophils, monocytes, and lymphocytes.

Vital signs (blood pressure and heart rate) and anthropometric data (weight and height) were collected upon admission. Laboratory parameters, including blood glucose, LDL, serum creatinine, uric acid, electrolytes (sodium and potassium), and NT-proBNP, were measured either at admission or from fasting blood samples collected on the following morning. Left ventricular ejection fraction (LVEF) was determined via echocardiography during hospitalization, and QRS duration was obtained from standard electrocardiograms recorded at admission.

#### 5.14.2 Statistical Analysis

Continuous variables were assessed for normality using the Shapiro-Wilk test. Normally distributed data are presented as mean ± standard deviation (SD) and were compared using the student’s t-test. Non-normally distributed variables are expressed as medians with interquartile ranges (IQR, 25th–75th percentiles) and were compared using the Mann-Whitney U test. Categorical variables are presented as frequencies and percentages, with comparisons performed using the chi-square test or Fisher’s exact test, as appropriate.

To minimize potential selection bias and balance baseline characteristics PSM was performed. The propensity score for each patient was calculated using a logistic regression model. The following variables were included in the PSM model as independent covariates: sex, age, smoking status, alcohol consumption, comorbidities (hypertension, diabetes, dyslipidemia, chronic kidney disease, angina pectoris, myocardial infarction, and atrial fibrillation), and baseline clinical parameters (NT-proBNP, QRS duration, serum creatinine, total cholesterol, LDL, hemoglobin, blood glucose, and uric acid). Medication use, including RAS inhibitors (ACEI/ARB/ARNI), diuretics, beta-blockers, and mineralocorticoid receptor antagonists (MRAs), was also accounted for.

Matching was performed using a nearest-neighbor algorithm with a caliper of 0.1. After PSM, the SGLT2i-treated group was matched with the non-treated group to ensure clinical comparability. For variables that remained unbalanced after PSM (e.g., NT-proBNP and diuretic use), stratified analyses were subsequently performed to validate the robustness of the hematological and inflammatory outcomes.

All statistical analyses were conducted using IBM SPSS Statistics version 22 (IBM Corp., Armonk, NY, USA). A two-sided (*P* < 0.05) was considered statistically significant.

## 6 List of abbreviations

Abbreviations: Full Name
SGLT2i: sodium glucose transport protein 2 inhibitors
EM: empagliflozin
DOCA: Deoxycorticosterone acetate
HSD: high salt diet
NT-proBNP: N-terminal pro-B-type natriuretic peptide
RAAS: Renin-angiotensin-aldosterone system
PVN: paraventricular nucleus of hypothalamus
CNS: central nervous system
IACUC: institutional animal care and use committee
SPRI: Solid Phase Reverse Immobilization
LVM: Left ventricular mass
LVEF: Left ventricular ejection fraction
TH: Tyrosine hydroxylase
NGS: Normal goat eerum
PFA: Paraformaldehyde
PBS: Phosphate buffered saline
DAPI: 4′,6-diamidino-2-phenylindole
ELISA: Enzyme linked immunosorbent assay
NE: Norepinephrine
IFN-γ: interferon γ
ACE2: Angiotensin converting enzyme 2
WB: Western blotting
HRP: Horseradish peroxidase
GAPDH: Glyceraldehyde-3-phosphate dehydrogenase
ECL: Enhanced chemiluminescence
PSM: Propensity score matching
HCT: Hematocrit
LDL: Low density lipoprotein
SD: Standard deviation
ARNI: Angiotensin receptor neprilysin inhibitor
ARB: Angiotensin II receptor blocker
ACEI: Angiotensin converting enzyme inhibitor
Hyd: Hydralazine
LV: Left ventricle
Cr: Creatinine
PCA: Principal component analysis
UMAP: Uniform manifold approximation and projection
GO: Gene ontology
DEG: Differentially expressed gene
AVP: Arginine vasopressin

## 7 Declarations

### 7.1 Institutional Review Board Statement

All procedures of anmial research were approved by the IACUC of Xi’an Jiaotong University (XJTU2021-203), in agreement with the Guide for the Care and Use of Laboratory Animals. Every effort was made to minimize the number of animals used and their suffering. Institutional Animal Care and Use Committees in accordance with the ‘Principles of Laboratory Animal Care by the National Society for Medical research and the Guide for the Care and Use of Laboratory Animals’ (National Institutes of Health Publication No. 86-23, revised 1996).

The clinical study of this study is a retrospective single-center clinical study. All participants provided written informed consent before participating in the study. The study was approved by the Institutional Review Board at the Second Affiliated Hospital of Xi’an Jiaotong University (IRB#2024-150). All methods were performed in accordance with the relevant guidelines and regulations.

### 7.2 Consent for publication

We confirm that the manuscript has been read and approved by all authors and that there are no other persons who satisfied the criteria for authorship but are not listed. We further confirm that the order of authors listed in the manuscript has been approved by all of us.

### 7.3 Availability of data and materials

All requests for raw data, analysed data and materials of this study will be promptly reviewed by the corresponding authors (Dengfeng Gao) and the Second Affiliated Hospital of Xi’an Jiaotong University.

The raw data of snRNA-seq are available in SRA (SUB15305597). The processed single-cell raw expression matrices that support the findings of this study are available on the Gene Expression Omnibus (GSE313832) at the following URL: https://identifiers.org/geo:GSE313832.

### 7.6 Authors’ contributions

DF.G., JX.X. and M.Y. (First author) are responsible for the conception of the article, the design of animal experiments and clinical research. DF.G. and JX.X. were responsible for the methodology and supervision of this study. M.Y. was responsible for the data analysis of single cell sequencing. M.Y., HX.W., JW.W. and ZH.Q. were responsible for the construction, collection and analysis of animal models. M.Y. and KX.L. were responsible for the collection and analysis of clinical data. M.Y. was responsible for visualization and wrote the original draft, which was revised by all the authors. All authors read and approved the final version of the manuscript.

## 7.7 Acknowledgements

We would like to thank Shenglan Yuan for her guidance and assistance in the animal experiment section. Thanks to the medical staff of the Department of Cardiovascular Medicine of the Second Affiliated Hospital of Xìan Jiaotong University for their help in the clinical research part. Thanks to all the patients who participated in the clinical research.

## Supplementary Materials

1. sup fig 1.jpg

**Supplementary Figure 1. Body weight, water consumption, and blood pressure profiles across experimental groups.**

A–B) Body weight curves of mice throughout the treatment period in the DOCA-salt (A) and high-salt diet (HSD) (B) models. (C–F) Daily water consumption curves (C, E) and corresponding area under the curve (AUC) statistical analysis (D, F) for the DOCA-salt (C, D) and HSD (E, F) models. (G–J) Statistical results of baseline systolic blood pressure (SBP) and diastolic blood pressure (DBP) measured before modeling in the DOCA-salt (G, H) and HSD (I, J) models. (K–N) Statistical results of final SBP and DBP measured at the end of the treatment period in the DOCA-salt (K, L) and HSD (M, N) models. Data are expressed as mean ± SEM (n = 10 per group). Statistical analysis was performed using one-way ANOVA followed by Tukey’s post-hoc test. \**P* < 0.05, \*\**P* < 0.01, \*\*\**P* < 0.001, \*\*\*\**P* < 0.0001; ns, not significant. Abbreviations: AUC, area under the curve; DBP, diastolic blood pressure; DOCA, deoxycorticosterone acetate; EM, empagliflozin; HSD, high-salt diet; Hyd, hydralazine; NSD, normal salt diet; SBP, systolic blood pressure.

2. sup fig 2.jpg

**Supplementary Figure 2. Pathological assessment of cardiac hypertrophy and fibrosis in salt-overload models.**

(A) Representative images of cardiac sections from the DOCA-salt model groups stained with Hematoxylin and Eosin (H&E) for general morphology (left), and Masson’s trichrome for interstitial fibrosis (middle) and perivascular fibrosis (right, focused on small vessels). (B–C) Statistical quantification of the collagen volume fraction (CVF) representing interstitial fibrosis (B) and perivascular fibrosis (C) in the left ventricle of the DOCA-salt model groups. (D) Representative H&E and Masson’s trichrome stained sections from the high-salt diet (HSD) model groups, showing interstitial and perivascular collagen deposition. (E–F) Statistical quantification of the CVF for interstitial (E) and perivascular (F) fibrosis in the HSD model groups. Data are presented as mean ± SEM (n = 5 per group). Statistical analysis was performed using one-way ANOVA followed by Tukey’s post-hoc test. \**P* < 0.05, \*\**P* < 0.01, \*\*\**P* < 0.001, \*\*\*\**P* < 0.0001; ns, not significant. Scale bars = 100 μm. Abbreviations: CVF, collagen volume fraction; DOCA, deoxycorticosterone acetate; EM, empagliflozin; H&E, hematoxylin and eosin; HSD, high-salt diet; Hyd, hydralazine; NSD, normal salt diet.

3. sup fig 3.jpg

**Supplementary Figure 3. Plasma osmolality, electrolytes, and hematological profiles across experimental groups.**

(A–B) Plasma osmolality in the DOCA-salt (A) and high-salt diet (HSD) (B) model groups, showing no significant systemic alterations across treatments. (C–D) Plasma sodium concentration in the DOCA-salt (C) and HSD (D) models. (E–F) Hematocrit (HCT) values measured from peripheral blood in the DOCA-salt (E) and HSD (F) models. Note that EM restored the DOCA-induced reduction in HCT. (G–H) Peripheral neutrophil counts determined by blood cell analysis in the DOCA-salt (G) and HSD (H) model groups. Data are presented as mean ± SEM (n = 5 per group). Statistical analysis was performed using one-way ANOVA followed by Tukey’s post-hoc test.\**P* < 0.05, \*\**P* < 0.01, \*\*\**P* < 0.001; ns, not significant. Abbreviations: DOCA, deoxycorticosterone acetate; EM, empagliflozin; HCT, hematocrit; HSD, high-salt diet; Hyd, hydralazine; NSD, normal salt diet.

4. sup fig 4.jpg

**Supplementary figure 4. Mapping of representative markers for the adjacent regions of PVN.**

5. Sup fig 5.jpg

**WB original image.**

6. sup tab 1.docx

**Supplementary Table 1. Comparison of HCT and peripheral immune cell counts between the treated group and non-treated group before propensity matching score.**

---

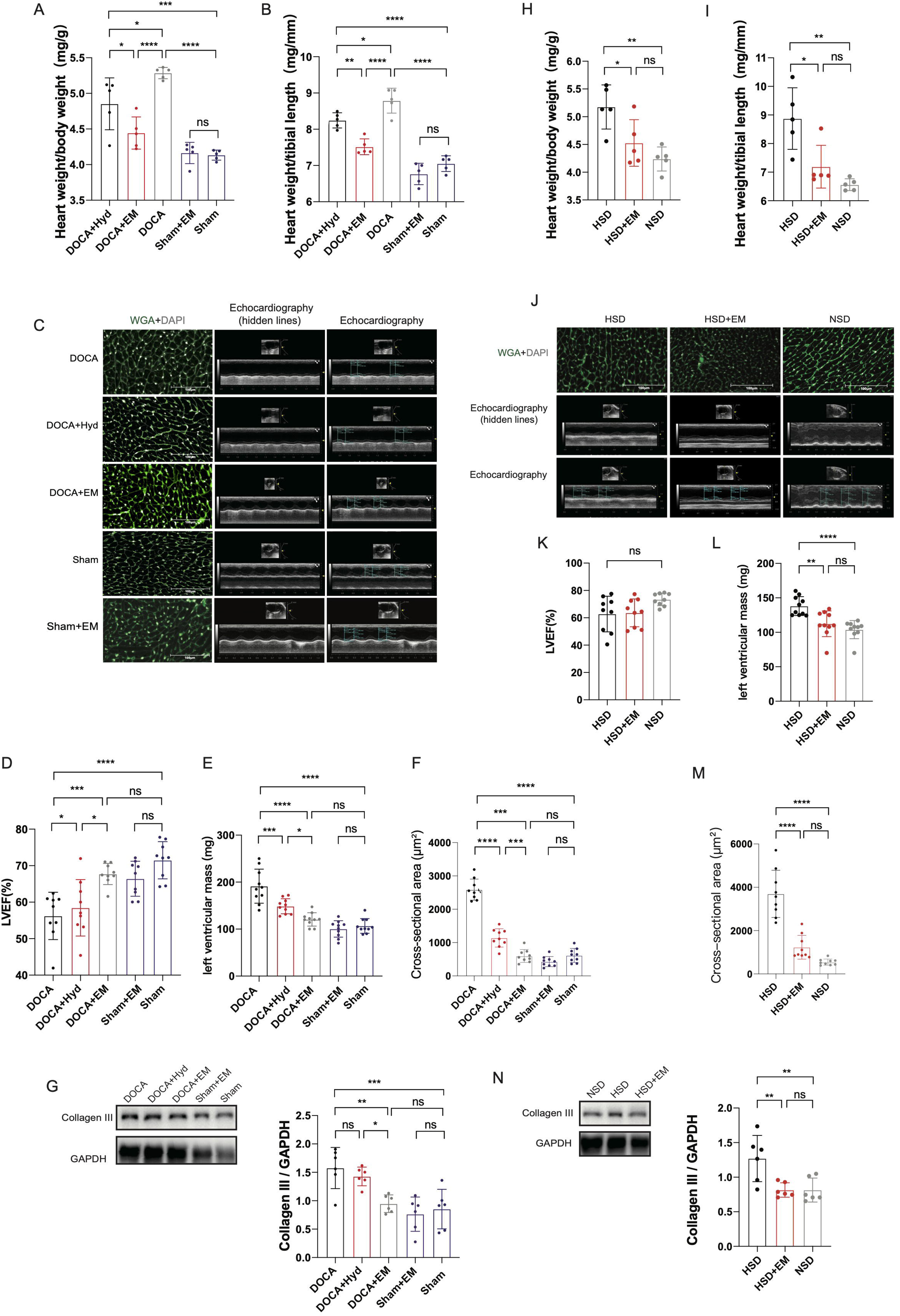

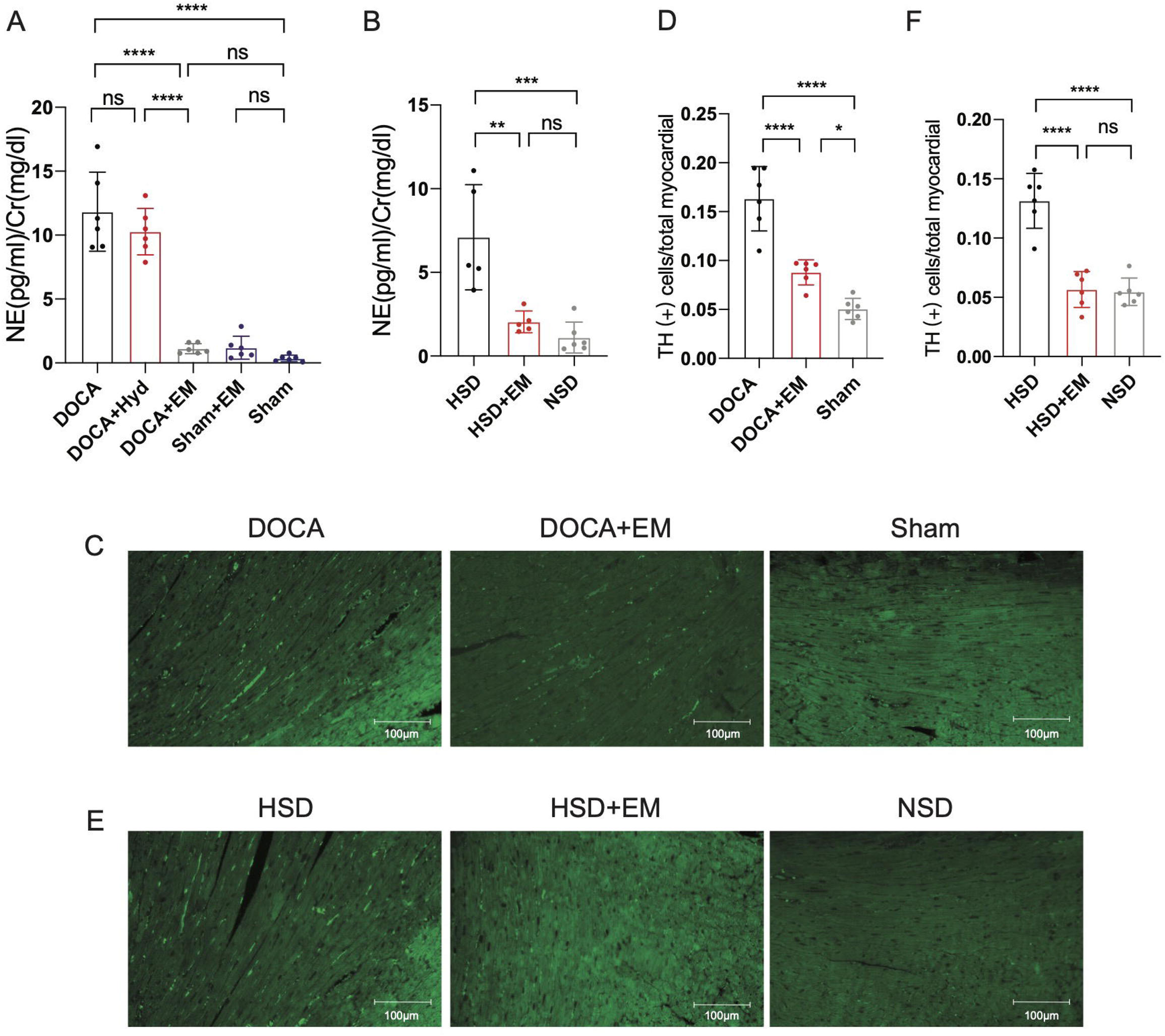

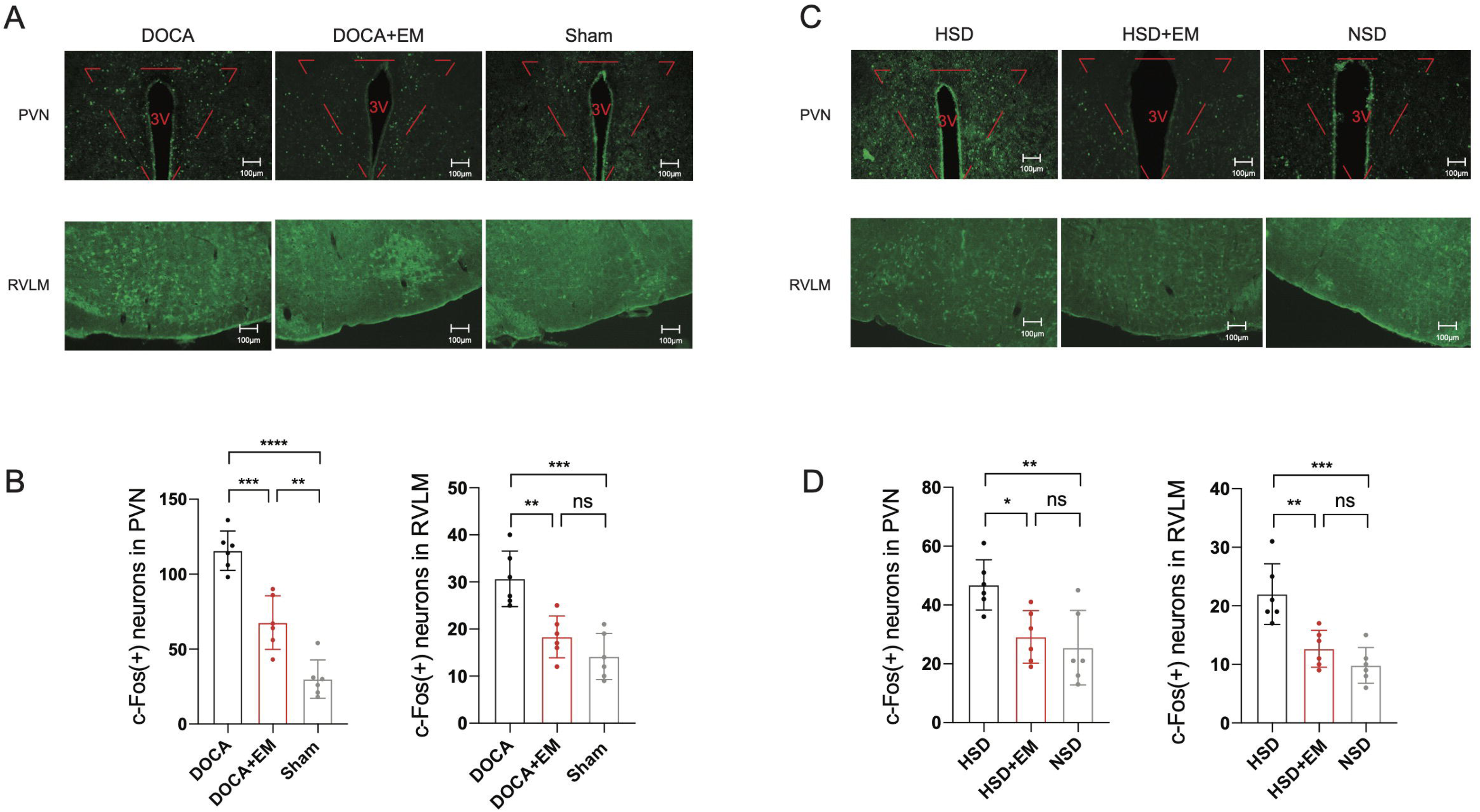

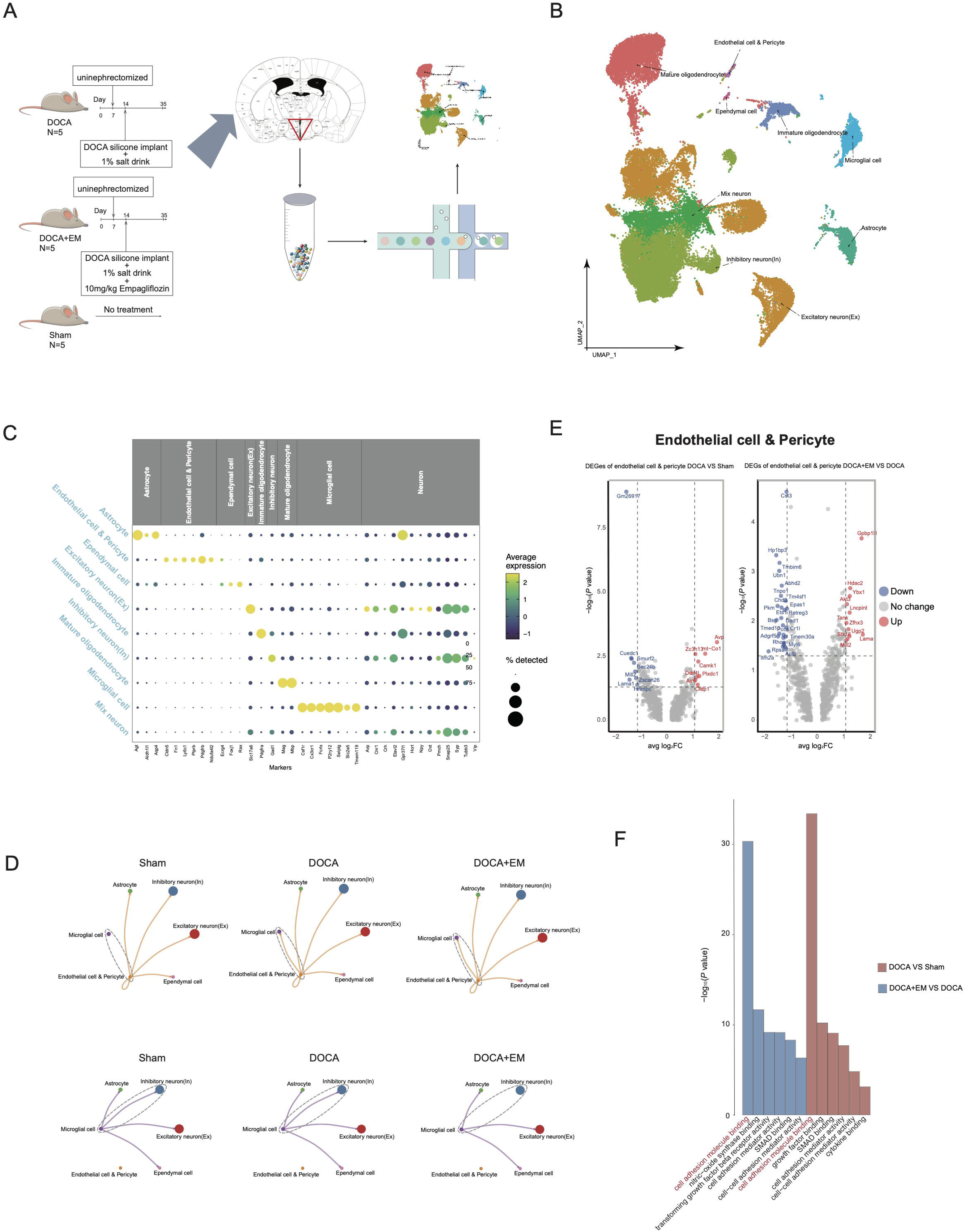

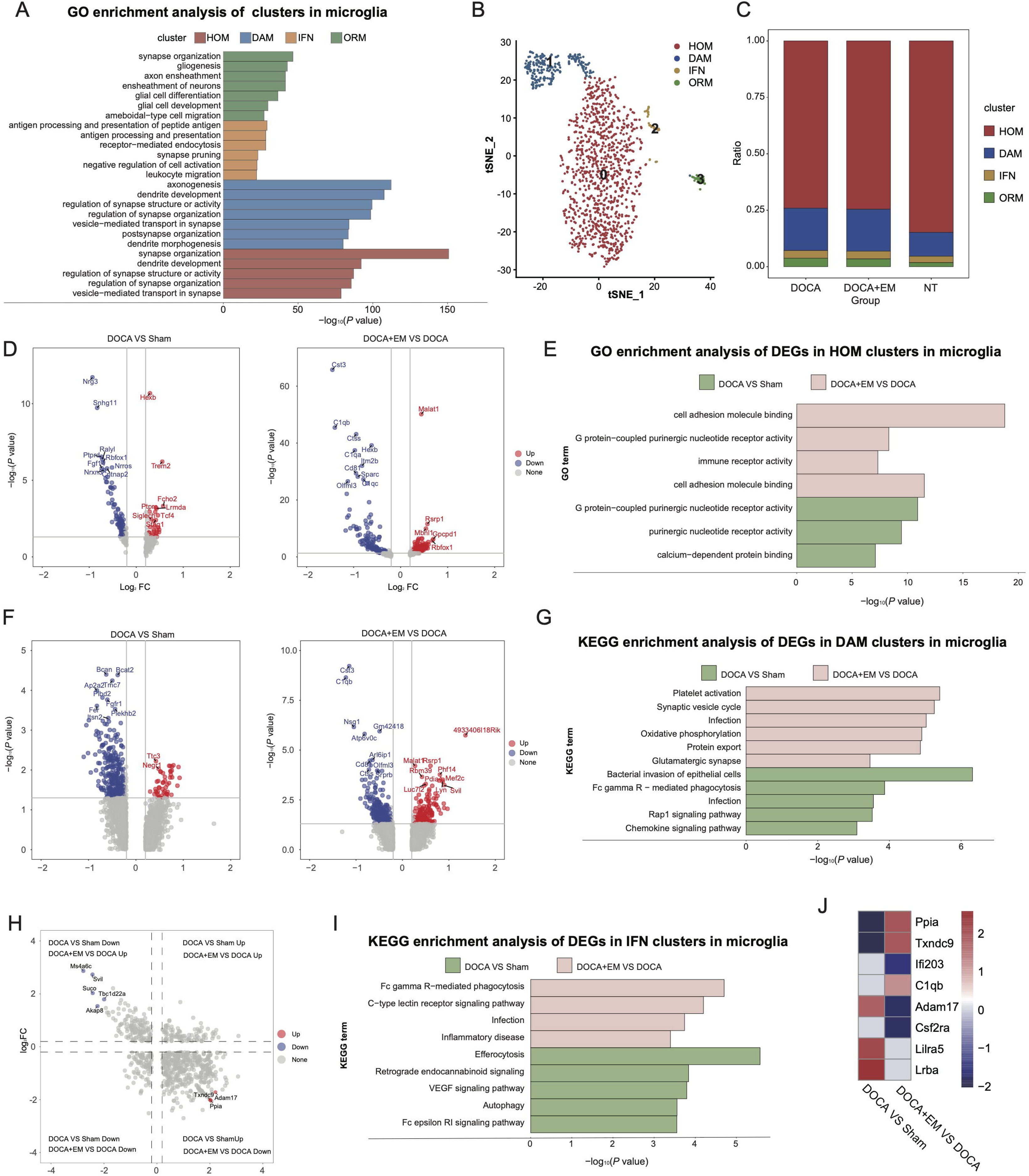

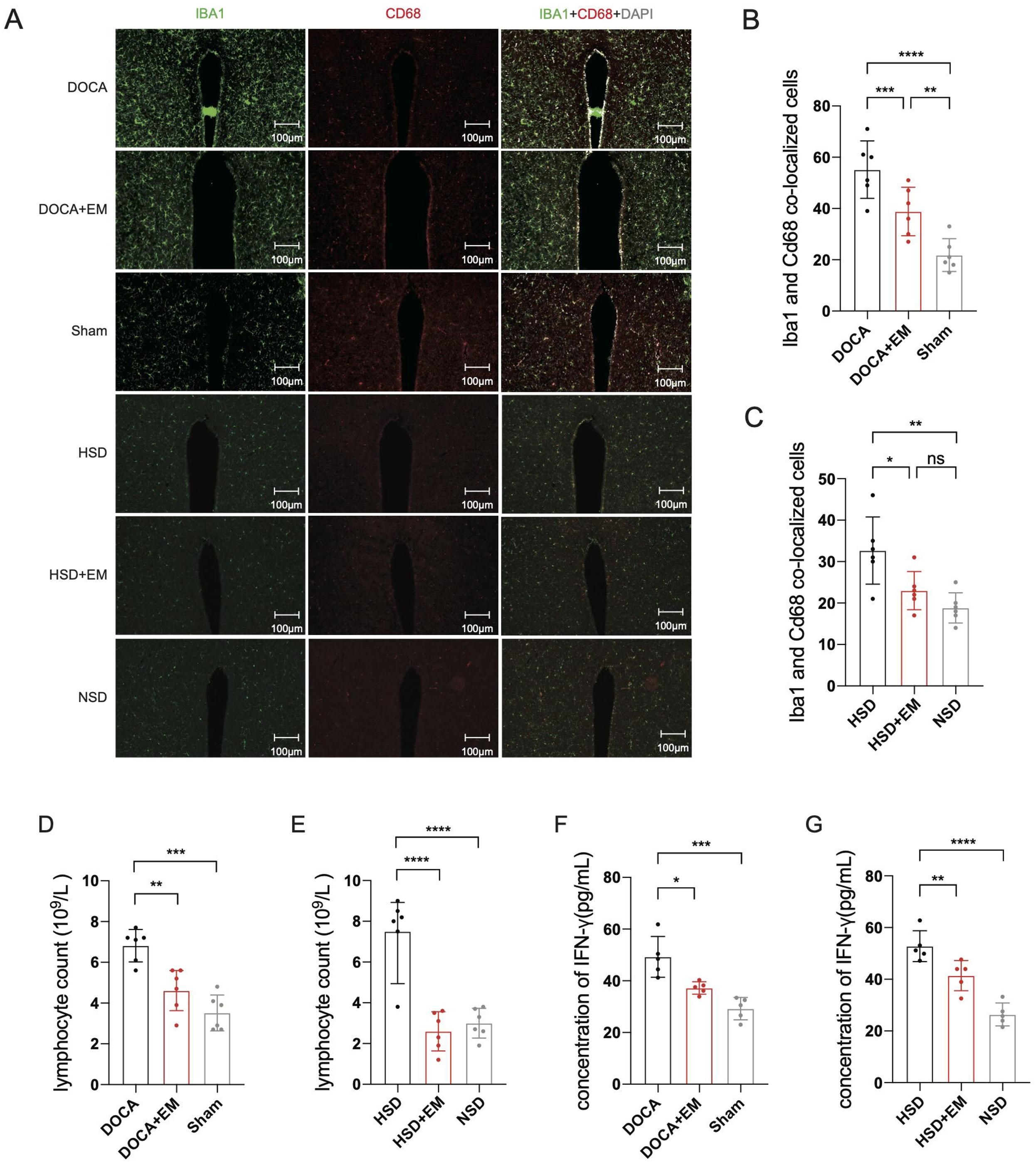

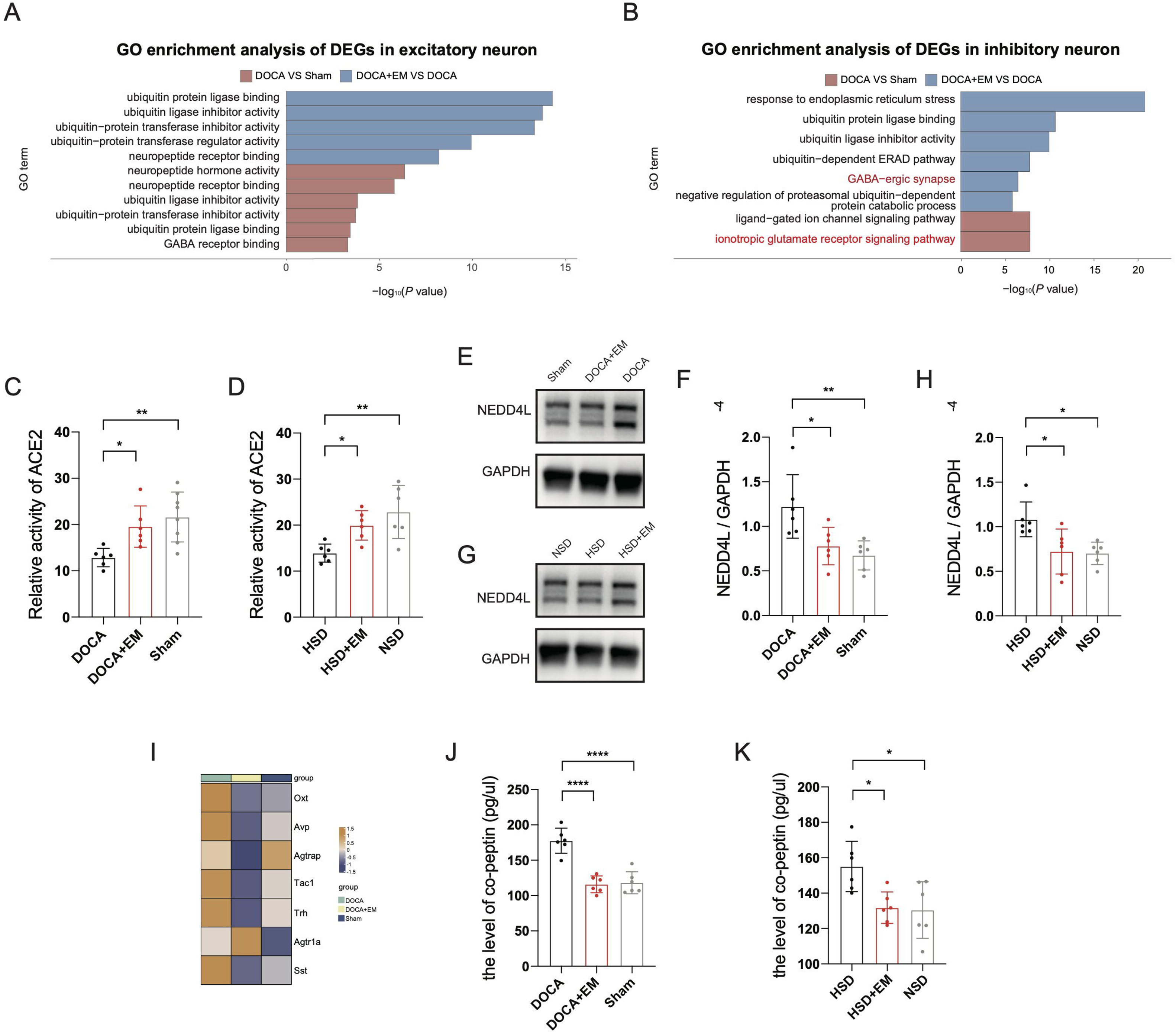

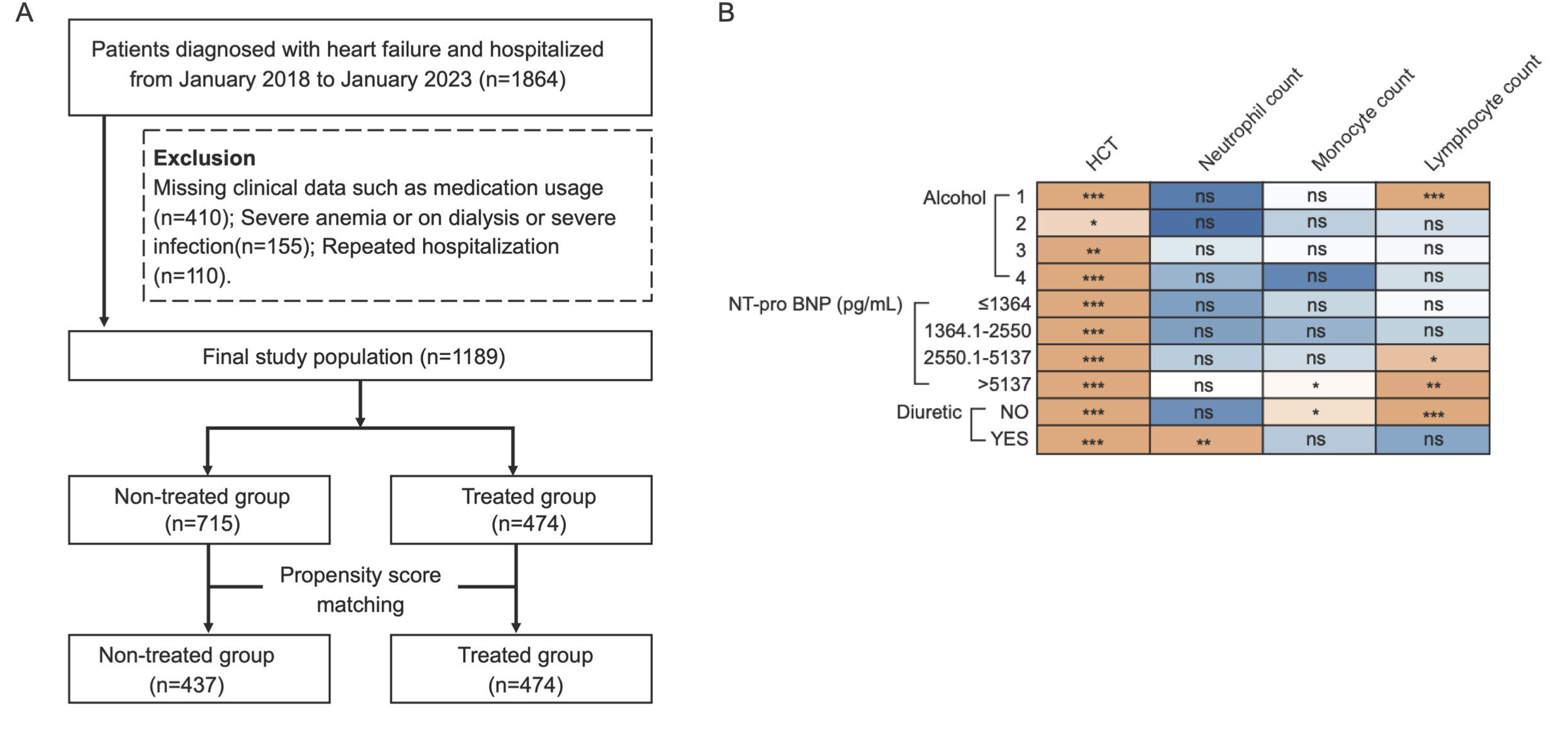

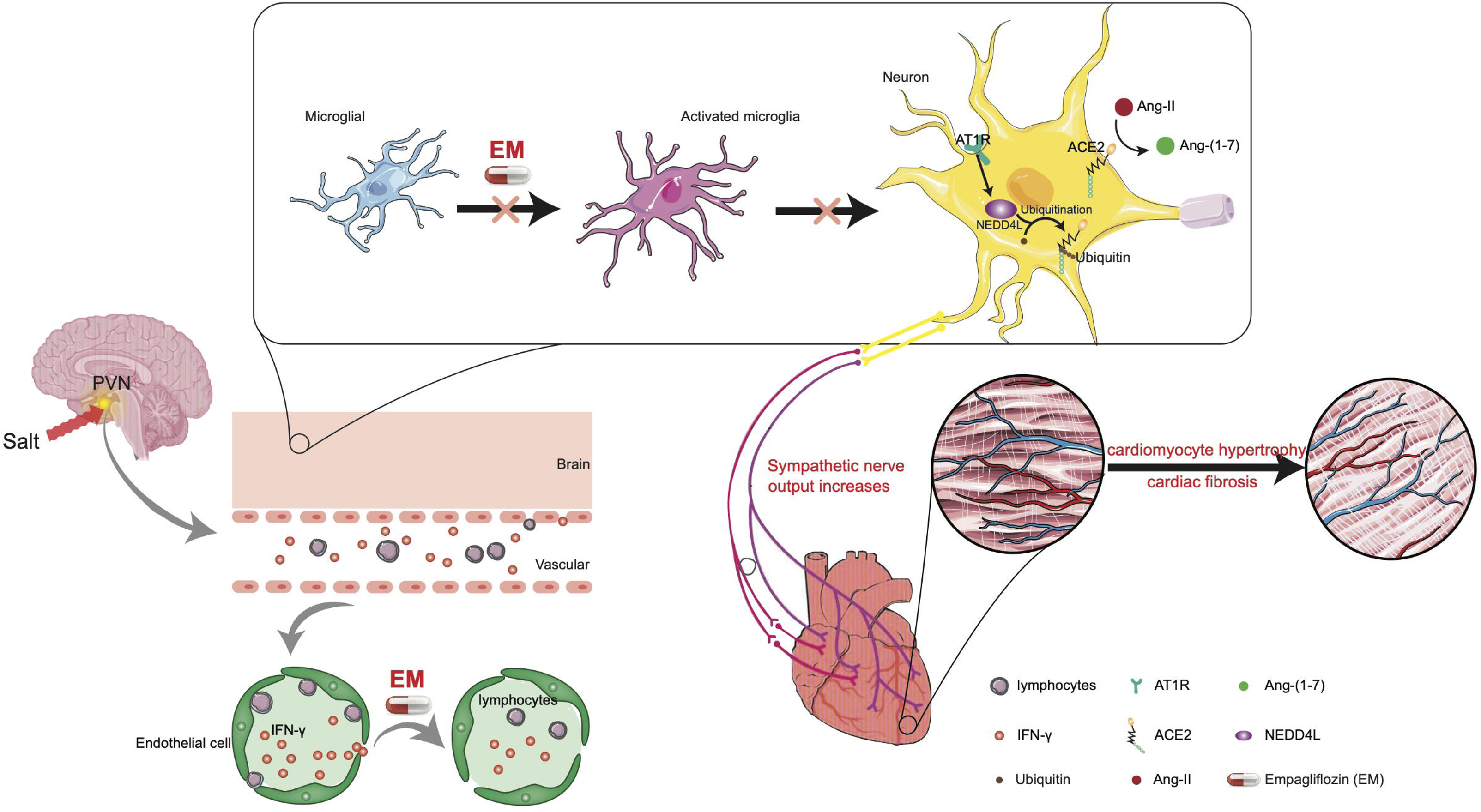

