## Supplementary figures and images for "Integrated analysis reveals neuro-immune pathway in the central nervous system that supports SGLT2i’s protective effects in treatment of cardiac remodeling"

### Supplementary Figure 1

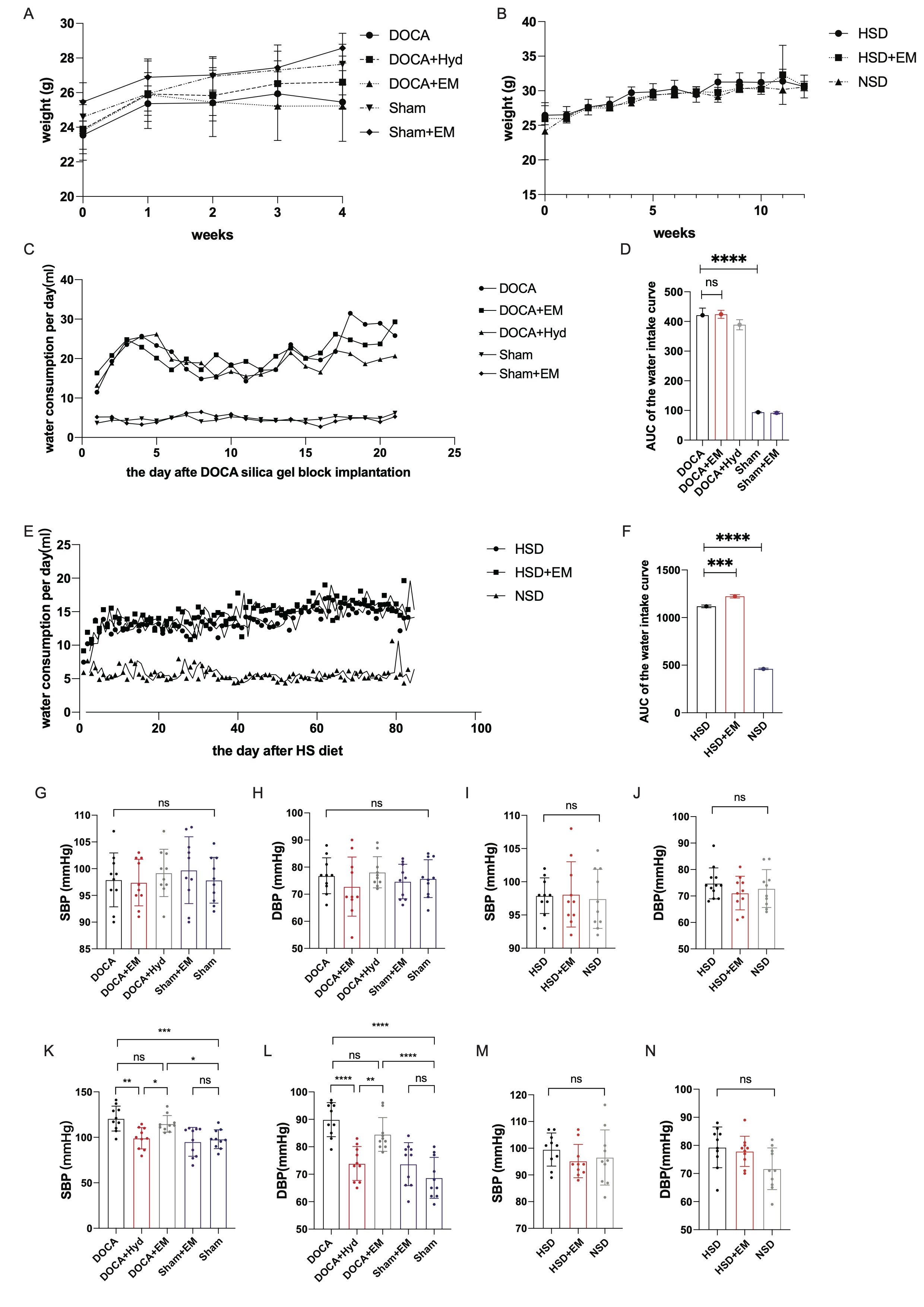

### Supplementary Figure 2

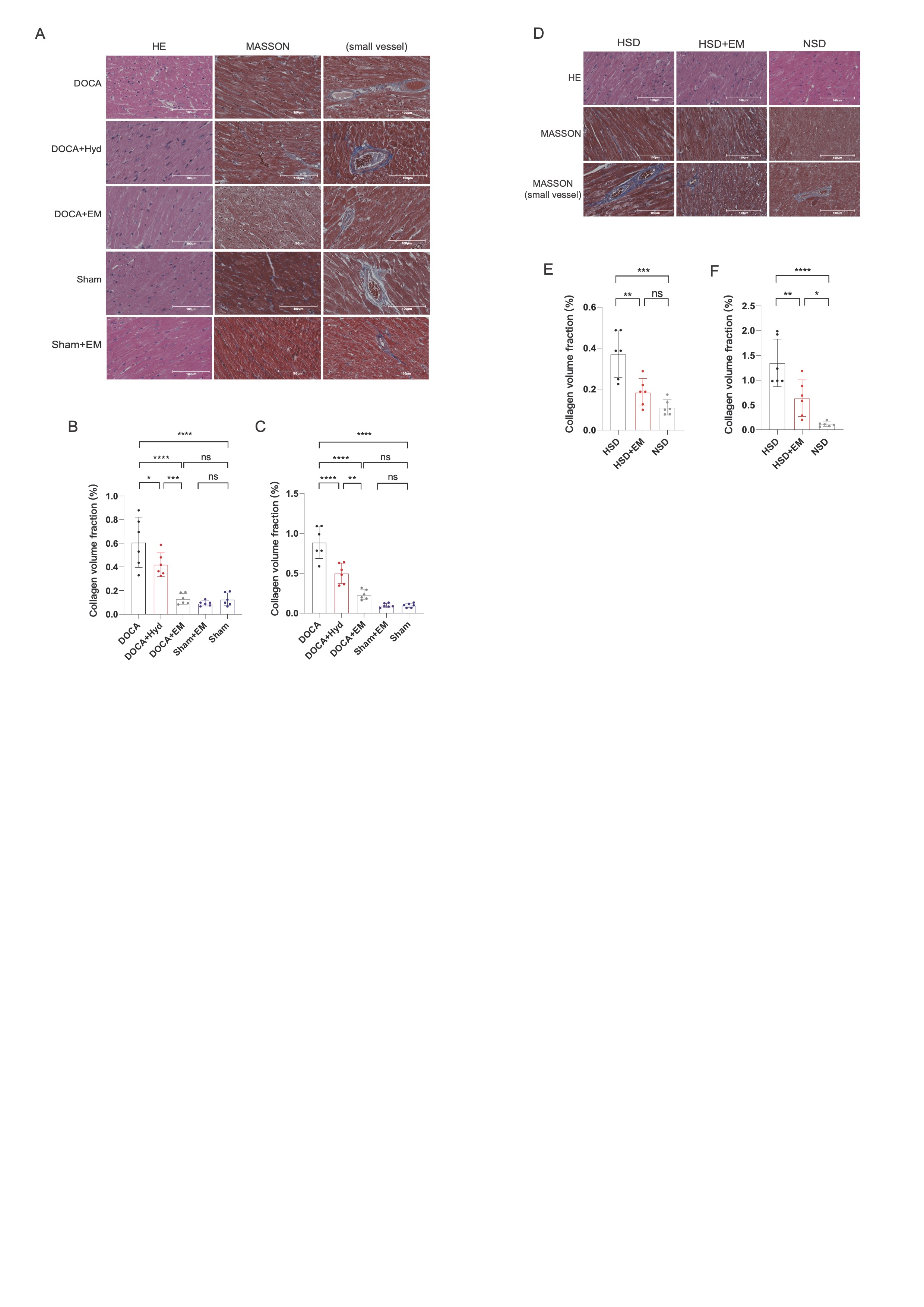

### Supplementary Figure 3

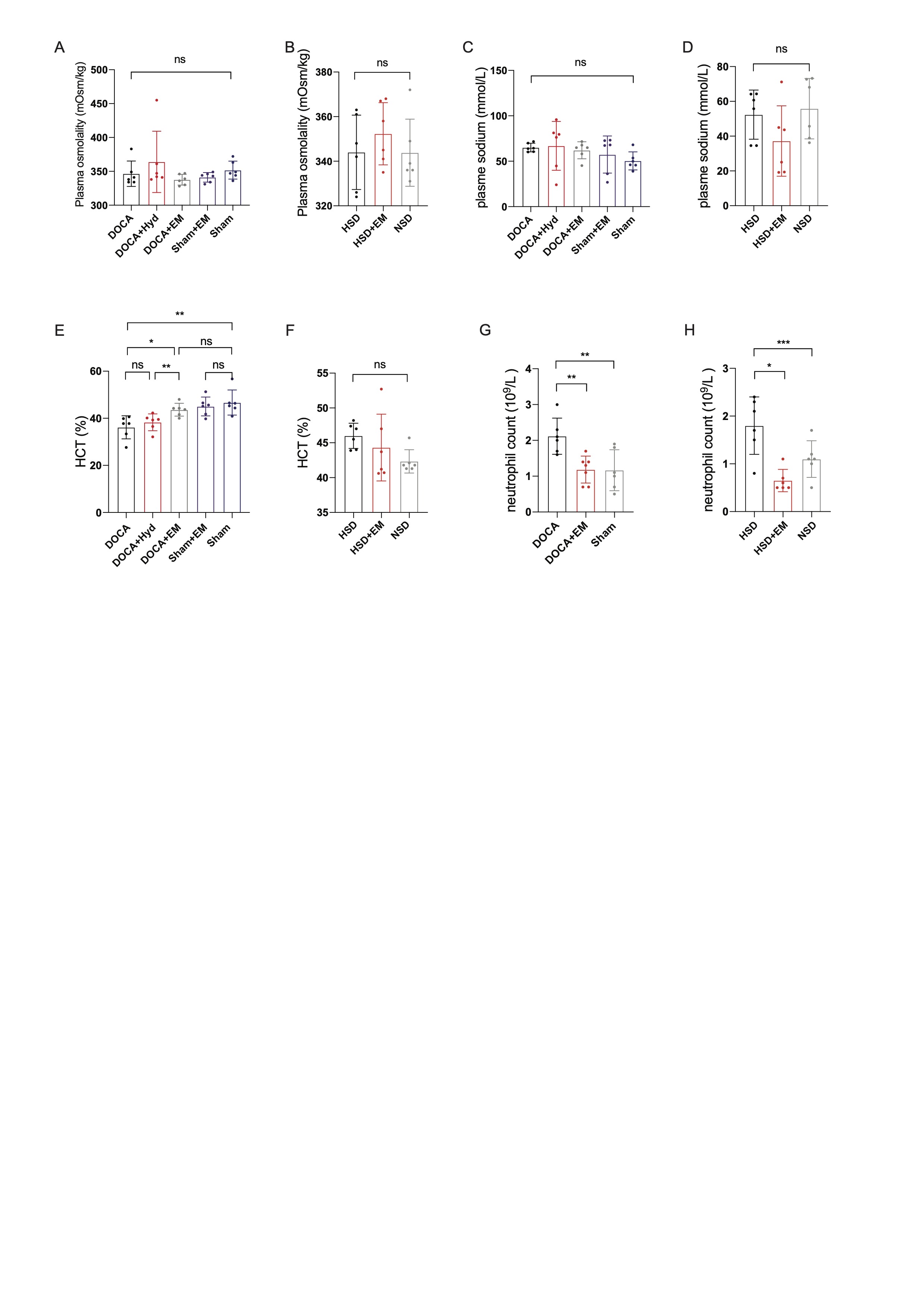

### Supplementary Figure 4

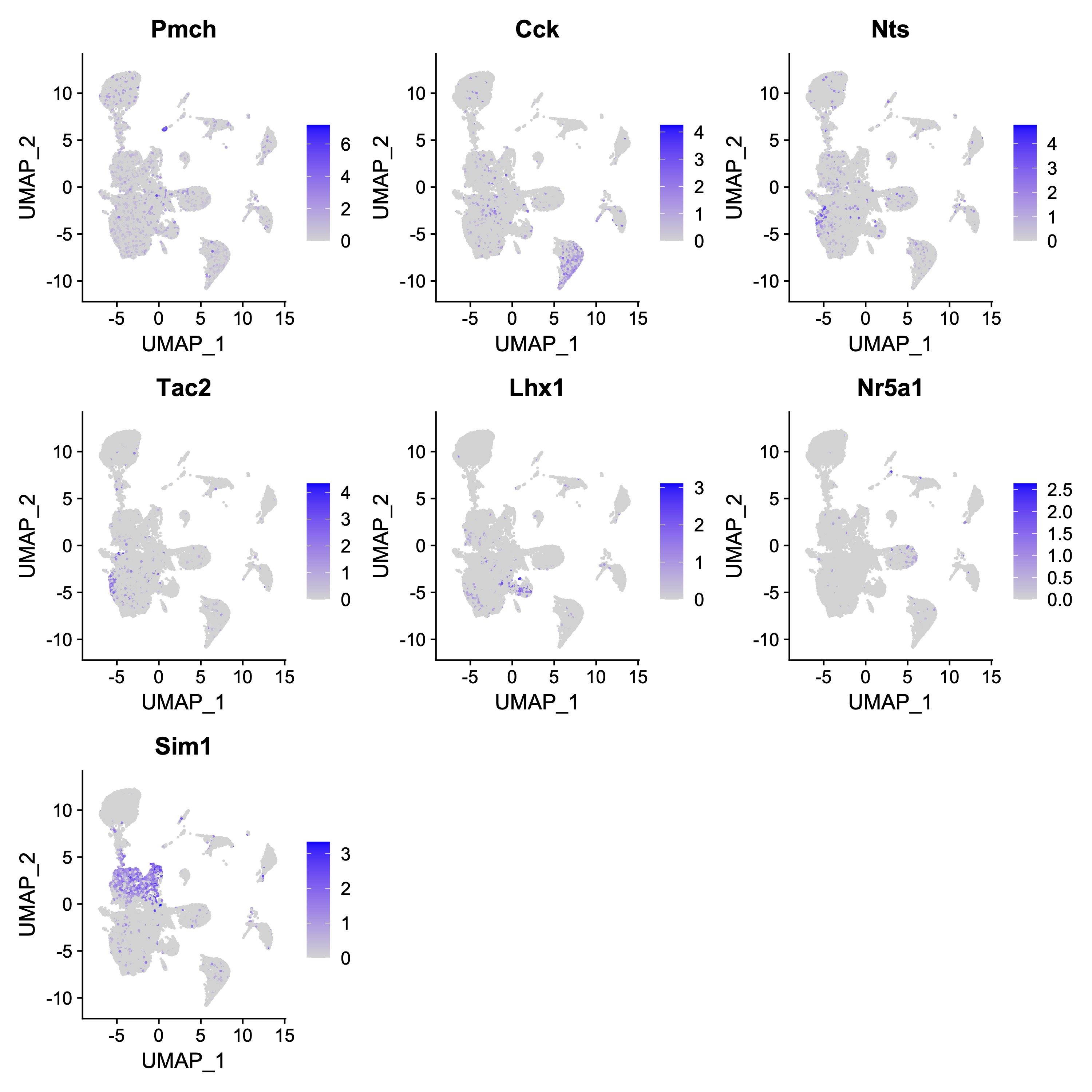

### Supplementary Figure 5

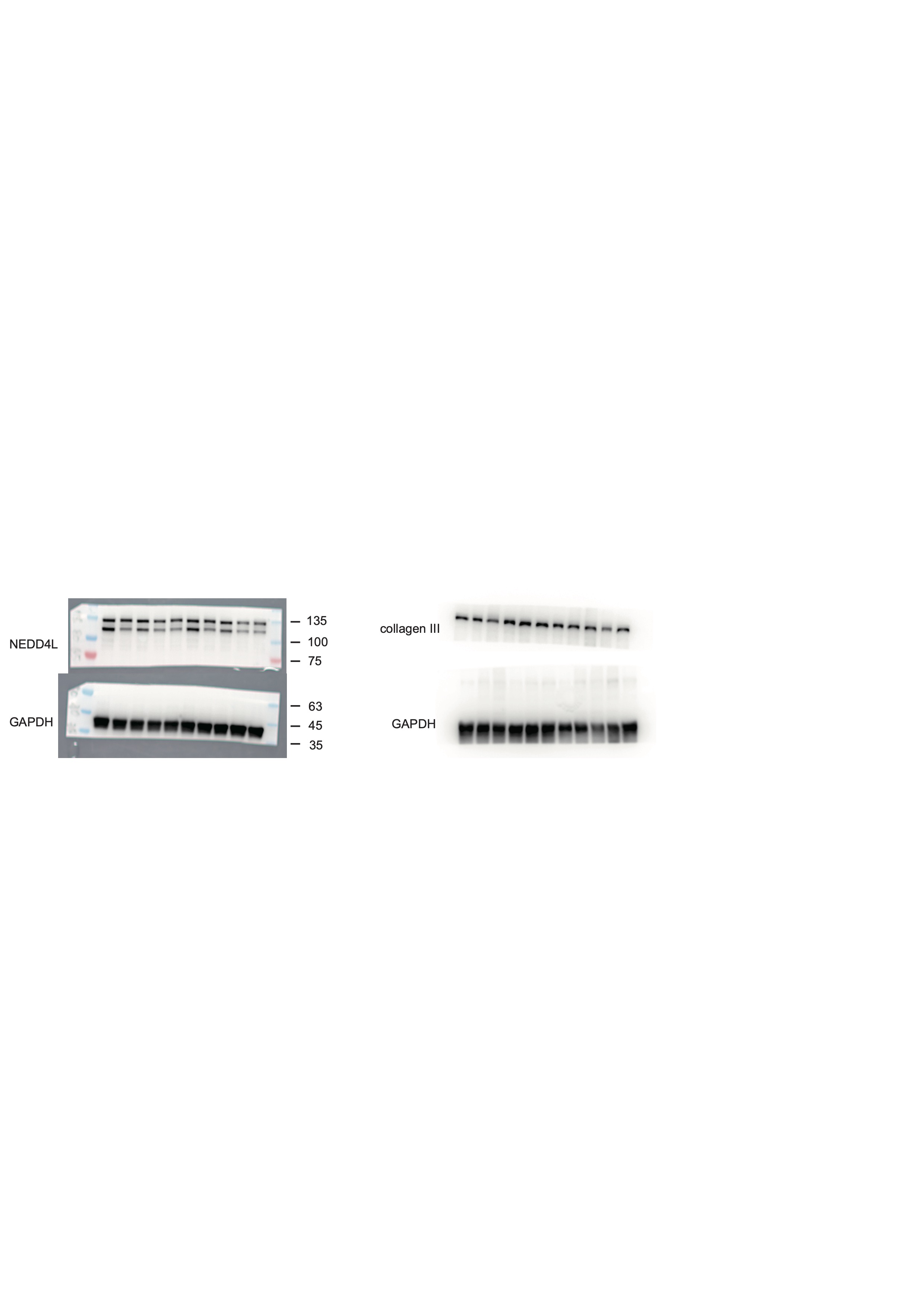
