## Supplementary Table 1 for "Integrated analysis reveals neuro-immune pathway in the central nervous system that supports SGLT2i’s protective effects in treatment of cardiac remodeling"

Supplementary Table 1 Comparison of HCT and peripheral immune cell counts between the treated group and non-treated group before propensity matching score

|  |  | Non-treat group | treat group | *P* value |
| --- | --- | --- | --- | --- |
|  | n=1189 | n=715 (60.1%) | n=474 (39.9%) |  |
| HCT |  | 39.5(36.5-43.2) | 43(39.63-46.1) | <0.001 |
| Neutrophil count |  | 4.16(3.18-5.43) | 4.18(3.19-5.31) | 0.99 |
| Monocyte count |  | 0.51(0.39-0.66) | 0.49(0.39-0.63) | 0.08 |
| Lymphocyte count |  | 1.5(1.19-1.9) | 1.39(1.0-1.77) | <0.001 |
